# Antisense lncRNA transcription drives stochastic Protocadherin α promoter choice

**DOI:** 10.1101/360206

**Authors:** Daniele Canzio, Chiamaka L. Nwakeze, Adan Horta, Sandy M. Rajkumar, Eliot L. Coffey, Erin E. Duffy, Rachel Duffié, Matthew D. Simon, Stavros Lomvardas, Tom Maniatis

## Abstract

Stochastic and combinatorial activation of clustered Protocadherin (Pcdh) α, β, and γ gene promoters generates a cell-surface identity code in individual neurons that functions in neural circuit assembly. Here we show that Pcdhα promoter choice requires transcription of a long noncoding RNA (lncRNA) initiated from newly identified promoters located in the protein coding sequence of each Pcdhα exon. Antisense transcription of the lncRNA through the sense promoter results in its activation and in DNA demethylation of the binding sites for the CCCTC-binding protein, CTCF, located in close proximity to both sense and antisense promoters. Increased CTCF binding promotes the assembly of long-range DNA contacts between the activated promoter and a neuron-specific enhancer, thus locking in the epigenetic state of the stochastically chosen Pcdhα promoter. Examination of this hierarchical molecular mechanism in differentiating olfactory sensory neurons, suggests that antisense Pcdhα transcription is a key prerequisite for stochastic Pcdhα promoter choice *in vivo*.

## INTRODUCTION

During brain development, individual neurons differentiate into distinct functional cell types, they respond to a plethora of guidance molecules, and project into specific regions of the nervous system to form complex neural circuits. A key aspect of this process is the ability of neurites of individual neurons (axons and dendrites) to distinguish between themselves and neurites from other neurons (self *vs* non-self) (Grueber and Sagasti, 2010; Lefebvre et al., 2015; Zipursky and Grueber, 2013). This process is required to avoid clumping or synaptic engagement between self-neurites, and at the same time, establish functional synaptic connections with other neurons of the same or different neuronal cell type (Grueber and Sagasti, 2010; Lefebvre et al., 2015; Zipursky and Grueber, 2013). The process by which neurites from the same neuron recognize and repel each other is known as self-avoidance and requires a unique combination of cell-surface homophilic recognition molecules that function as a molecular identity code or barcode (Zipursky and Grueber, 2013; Zipursky and Sanes, 2010). Previous studies have identified the nature and function of such barcodes in invertebrates and vertebrates (Mountoufaris et al., 2017; Zipursky and Grueber, 2013; Zipursky and Sanes, 2010). In flies, the Drosophila down syndrome cell adhesion molecule (Dscam1) provides unique cell surface barcodes for neuronal self-avoidance (Hattori et al., 2008). In this case, stochastic alternative RNA splicing of a Dscam1 precursor messenger RNA (pre-mRNA) in individual neurons can generate over 19,000 distinct extracellular protein isoforms (Wojtowicz et al., 2004). In an extraordinary example of convergent evolution, the same cell-surface mechanism involving specific homophilic interactions followed by repulsion is used for self-avoidance in vertebrates. However, in this case, protein diversity is provided by the clustered Protocadherins (Pcdhs) rather than by Dscam1 homolog proteins (Chen et al., 2013; Mountoufaris et al., 2017; Yagi, 2013; Zipursky and Sanes, 2010). Moreover, rather than stochastic alternative RNA splicing of a single Dscam1 pre-mRNA in flies, the extraordinary functional diversity of clustered Pcdhs is a consequence, at least in part, of the unique genomic arrangement of the Pcdh genes in three closely linked clusters (designated as α, β, and γ) and a mechanism of stochastic and combinatorial promoter choice which remains poorly understood (Esumi et al., 2005; Tasic et al., 2002; Wang et al., 2002; Wu and Maniatis, 1999a; Wu et al., 2001).

The three Pcdh gene clusters, together, span nearly 1 million base pairs (bp) of DNA, and are organized into variable and constant regions, reminiscent of the organization of immunoglobin and T-cell receptor gene clusters (Wu and Maniatis, 1999b). The variable regions in the Pcdh α and γ cluster are further distinguished into alternate and c-types. The organization of the human Pcdh gene cluster, which is conserved throughout vertebrate evolution, is illustrated in Figure 1A. Neuron-specific expression of individual Pcdhα genes requires long-range DNA looping between random Pcdhα promoters and a transcriptional enhancer, called HS5-1 (hypersensitivity site 5-1) (Guo et al., 2012; 2015; Kehayova et al., 2011; Monahan et al., 2012; Ribich et al., 2006) (Figure 1B). Conserved transcriptional promoter sequences are located immediately proximal to every Pcdhα exon (Tasic et al., 2002) while the HS5-1 enhancer is located downstream of the constant exons, between the Pcdh α and the β clusters (Ribich et al., 2006) (Figure 1A). These stochastic promoter/enhancer interactions occur independently on each of the two allelic chromosomes in diploid cells and require the binding of the CCCTC-binding protein (CTCF) and the Cohesin protein complex (Guo et al., 2012; Hirayama et al., 2012; Kehayova et al., 2011; Monahan et al., 2012) (Figure 1B and 1C). CTCF is an 11-zinc finger domain protein that, together with the Cohesin complex, plays a central role as an insulator of chromatin domains, and mediates genome-wide promoter/enhancer interactions (Carretero et al., 2010; Ghirlando and Felsenfeld, 2016; Ong and Corces, 2014). All Pcdhα alternate exons contain two CTCF binding sites (CBS), one in the promoter (pCBS) and the other in the protein coding sequence in the downstream exon (eCBS) (Guo et al., 2012; Monahan et al., 2012) (Figure 1B). The two binding sites are separated by approximately 1000 base pairs, and similarly spaced CBS sites are located in the HS5-1 enhancer (L-CBS and R-CBS) (Guo et al., 2012; Monahan et al., 2012) (Figure 1B). Interestingly, the CTCF binding sites in Pcdhα promoters and the HS5-1 enhancer are in opposite relative orientations, and inversion of the HS5-1 enhancer results in a significant decrease in Pcdhα gene cluster expression, demonstrating the functional importance of this arrangement (Guo et al., 2015). This opposite relative orientation of promoter and enhancer CTCF binding sites appears to be a general feature of eukaryotic chromosomes genome-wide, suggesting that the relative orientation of CTCF binding sites plays an important role in determining the specificity of enhancer/promoter interactions (Guo et al., 2015; Rao et al., 2014).

**Figure 1:**
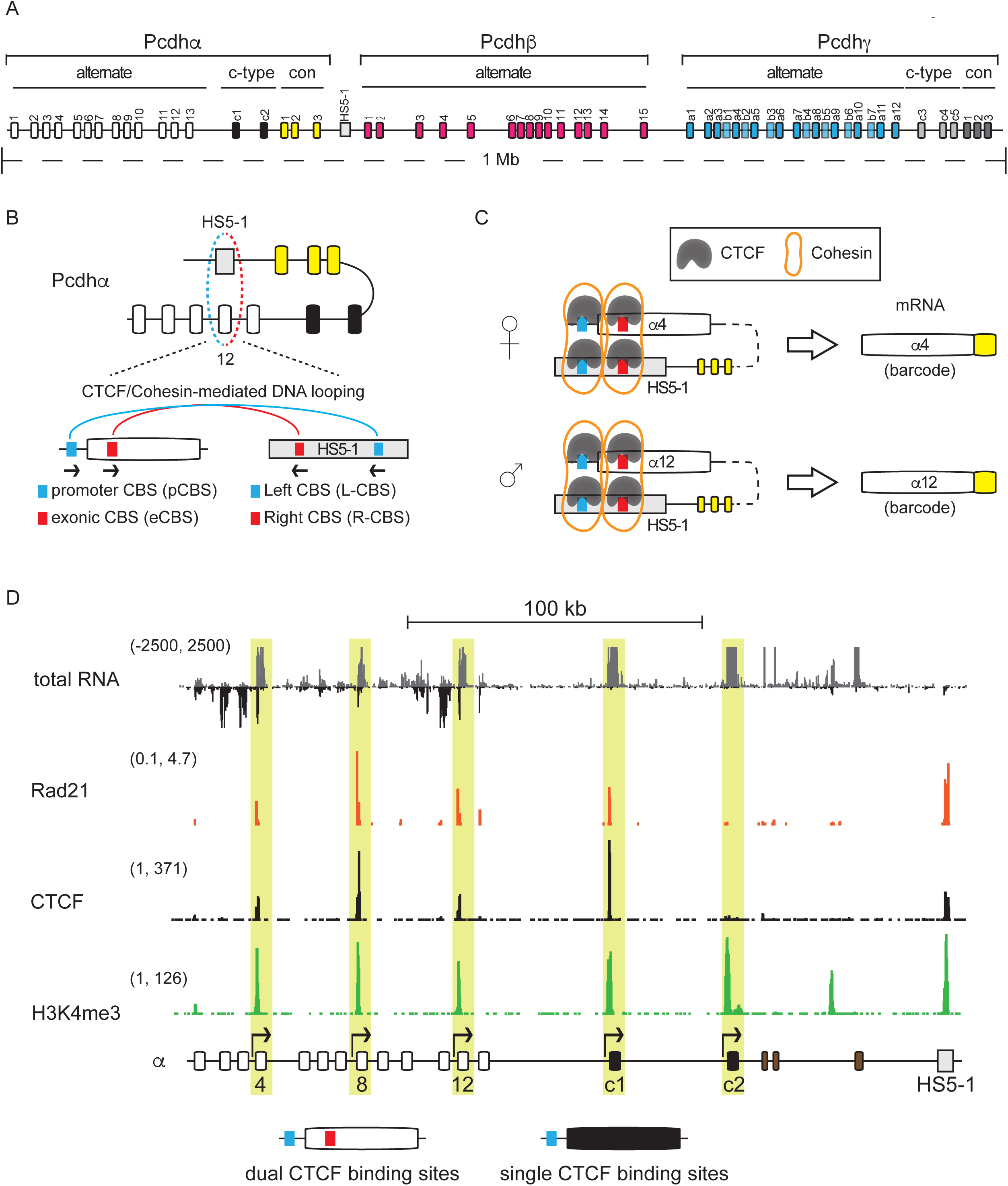
Transcription of sense and antisense RNA from Pcdhα alternative exons. (A) Genomic organization of the human Pcdh gene cluster. (B) Top: an example of DNA looping between the HS5-1 enhancer and a Pcdhα promoter (Pcdhα12 is shown). Bottom: Architecture of the CTCF binding sites (CBSs) in Pcdhα alternative exons and the HS5-1 enhancer. The CTCF/Cohesin complex mediates DNA looping between a chosen promoter and the HS5-1 enhancer. (C) Chromosome-independent promoter/enhancer long-range DNA looping. (D) Sense (grey) and antisense (black) transcription from the Pcdhα cluster in SK-N-SH cells. Distribution of CTCF, Rad21 (a Cohesin subunit) and H3K4me3 assayed by ChIP-Seq indicate transcriptionally active exons in SK-N-SH cells. Pcdhαc2 is active but not bound by CTCF or Rad21. The active exons are highlighted in yellow. The x-axis represents the linear sequence of the genomic organization of the Pcdhα cluster. The numbers on the left-hand side of each track represent the minimum and maximum densities in read per million.

An additional insight into the formation of a Pcdhα promoter/enhancer complex is provided by the observation that there is an inverse relationship between Pcdhα gene expression and DNA methylation of the GpC-rich CTCF binding sites in the Pcdhα promoters (Tasic et al., 2002; Toyoda et al., 2014). Specifically, the CTCF/Cohesin complex associates exclusively with active promoters, which are characterized by hypomethylation of the CTCF binding sites (CBS) (Guo et al., 2012). By contrast, CBS sites within inactive promoters are hypermethylated and appear refractive to CTCF/Cohesin binding (Guo et al., 2012). Yet, despite the fact that DNA methylation of the CTCF binding sites is likely to play an important role in the mechanism of stochastic Pcdhα promoter choice by the CTCF/Cohesin complex, the time during neuronal differentiation at which Pcdhα promoter choice is not known, nor is the ground state of Pcdhα promoter DNA methylation known. For instance, is the ground state of the Pcdhα gene cluster methylated and repressed, and its activation involves stochastic demethylation? Or, is the ground state unmethylated and promoter choice depends on stochastic activation followed by methylation of the promoters that are not chosen?

Here, we use a combination of cell-culture and *in vivo* model systems, to provide evidence that the ground state of the DNA of Pcdhα promoters is methylated and transcriptionally silent, and transcription of an antisense lncRNA is required to demethylate, derepress and activate Pcdhα mRNA transcription through a CTCF/Cohesin-dependent long-range DNA looping between the promoter and the Pcdhα neuronal-specific HS5-1 enhancer.

## RESULTS

### Transcription of sense and antisense RNA from clustered Pcdhα alternate exons

The formation of a promoter/enhancer-CTCF/Cohesin complex plays a critical role in the mechanism of stochastic promoter activation of Pcdhα alternate exons (Guo et al., 2015; Kehayova et al., 2011; Monahan et al., 2012; Ribich et al., 2006). However, the mechanism by which random Pcdhα promoters are activated is not understood and mechanistic studies of promoter choice are not possible *in vivo*, as every neuron expresses a distinct repertoire of Pcdhα alternate exons. Therefore, we made use of the well-characterized human neuroblastoma cell line SK-N-SH, which stably expresses a distinct repertoire of Pcdhα isoforms through multiple cell divisions: α4, α8, α12, αc1, and αc2 (Guo et al., 2012) (Figure 1D). This stochastic pattern of expression is indistinguishable from that observed in single neurons *in vivo* (Esumi et al., 2005; Mountoufaris et al., 2017). SK-N-SH cells thus provide a multicellular “avatar” for studying single cell expression of Pcdh genes and they provide internal controls for exons that are transcriptionally silent.

Another challenge to the study of Pcdhα promoter choice is the low level of expression of Pcdh genes. To optimize the analysis of Pcdh RNA precursors (pre-mRNA) and mature (mRNA) RNAs in SK-N-SH cells, we used capture RNA-Sequencing (cRNA-Seq) which allowed an enrichment for Pcdh RNA transcripts up to two orders of magnitude (Figure S1A and S1B). Remarkably, cRNA-Seq performed in SK-N-SH cells revealed a previously undetected high level of antisense RNA transcription originating within the transcriptionally active Pcdhα alternate exons (Pcdhα 4, 8 and 12) (Figure 1D and S1B). By contrast, antisense RNA transcription was not detected within the two c-type exons, αc1 and αc2, which do not contain CBSs within their exons (Figure 1D). Similarly, antisense RNA was not observed in the Pcdh β or γ variable exons in SK-N-SH cells, which also do not contain exonic CBS sites (Figure S1B). We refer to the antisense RNA as antisense long noncoding RNA, as-lncRNA, as it appears to lack protein-coding sequences, based on analyses of the open reading frames. For clarity, we refer to the sense Pcdh coding RNA as, s-cRNA (sense coding RNA).

The cRNA-Seq data obtained from SK-N-SH cells revealed a direct correlation between sense and antisense RNA transcription and transcriptionally active Pcdhα alternate exons (Figure 1D). Because transcription of the Pcdhα alternate exons occurs independently on the two allelic chromosomes (Esumi et al., 2005), we sought to determine whether the as-lncRNA and the s-cRNA were transcribed from the same allele. To address this, we used CRISPR-Cas9 gene editing and generate SK-N-SH cells heterozygous for the Pcdhα gene cluster (Figure S2, SK-N-SH-αhet). We isolated two clones (SK-N-SHαhet 1 and 2) expressing α12, αc1 and αc2 from the remaining copy of the Pcdhα gene cluster (Figure S3A). Both clones showed expression of the as-lncRNA and s-cRNA from Pcdhα12 (Figure S3A and S3B), confirming that sense and antisense transcription originate from the same allele. For one of the two clones isolated, αhet-1, we also performed chromatin immunoprecipitation studies (ChIP-Seq) for CTCF and Rad21, a subunit of the Cohesin complex, as well as *in situ* capture chromosome conformation studies (cHi-C) to examine long-range DNA interactions between the active Pcdhα12 and the HS5-1 enhancer (Figure S3A and S3C). These studies demonstrated that the Pcdhα alternate exons from which sense and antisense RNAs are transcribed are bound by the CTCF/Cohesin complex and engaged in a promoter-enhancer complex (Figure S3A). It is notable that the αhet-1 and αhet-2 clones share a 16.7 kilobase deletion that truncates the Pcdhα8 exon and removes the Pcdh α9 and α10 exons (Figure S3C). Remarkably, this deletion was previously reported as a common feature of individuals from multiple populations of European and East Asian descent with no apparent phenotypic consequence (Noonan, 2003). This deletion appears to affect both sense and antisense RNA expression from the Pcdhα8 exon (data not shown).

These data show that transcriptionally active Pcdhα alternative exons express both sense and antisense RNAs. Thus, in contrast to the SK-N-SH cells, a mixed population of primary neurons, each expressing a distinct repertoire of Pcdhα alternative exons, should express as-lncRNAs from the Pcdhα 1 to 13 exons, but not from Pcdh αc1 and αc2, or from the β or γ exons. As predicted, analysis of RNA from human primary neurons revealed lncRNA expression exclusively from the Pcdhα 1-13 exons, and from the HS5-1 enhancer (Figure S4A). Similarly, analysis of mouse olfactory sensory neurons (OSNs) also revealed lncRNA expression by all the Pcdhα alternate exons (Figure S4B). Thus, all Pcdhα alternative exons in human cell lines and human and mouse primary neurons, explored in this study, express as-lncRNAs. Furthermore, our RNA-Seq data revealed that the Pcdhα as-lncRNAs are polyadenylated (Figure S4).

### Convergent promoters in the Pcdhα alternative exons and HS5-1 enhancer

In order to characterize the nature of the antisense RNAs and to explore their function, we localized the promoters driving their transcription by identifying the location of promoter-paused RNAPII using Start-Seq (Nechaev et al., 2010) (Figure 2A). RNA isolated from stalled RNAPII at promoters are approximately 15 to 45 nucleotides long, and contain a 5’ 7meG-cap (Figure 2A). Isolation and 3’ end sequencing of these short RNAs reveals the position of paused RNAPII, and thus acts as a proxy for the location of RNAPII-engaged promoters (Figure 2A). As expected, we observed RNAPII pausing at the pCBS-promoter of the active Pcdh α4, α8, α12 and αc1 exons, and at the promoter of αc2 in SK-N-SH cells (Figure 2B, Pcdh α4 and αc1 are shown). However, we also observed RNAPII pausing just upstream of the eCBS for α4, α8, and α12 in the antisense orientation (Figure 2B, Pcdhα4 is shown). Thus, the two CBSs in the active Pcdhα genes act as convergent promoters (Figure 2C). This is in contrast to the sole pCBS site in Pcdhαc1, which acts as a divergent promoter (Figure 2C). Start-Seq analysis also identified a similar convergent promoter architecture of the two CBSs in the HS5-1 enhancer (Figure 2B and 2C). Mapping the location of the as-lncRNA promoters with respect to the as-lncRNAs revealed that the nuclear RNA precursors can be as long as 20,000 base pairs (Figure 1D). As an example, the as-lncRNA that initiates at the eCBS-promoter of Pcdhα4 in SK-N-SH cells is transcribed through the pCBS-promoter of Pcdhα4 and extends in the antisense direction all the way to the Pcdhα1 exon (more than 20,000 nucleotides upstream) (Figure 1D).

**Figure 2:**
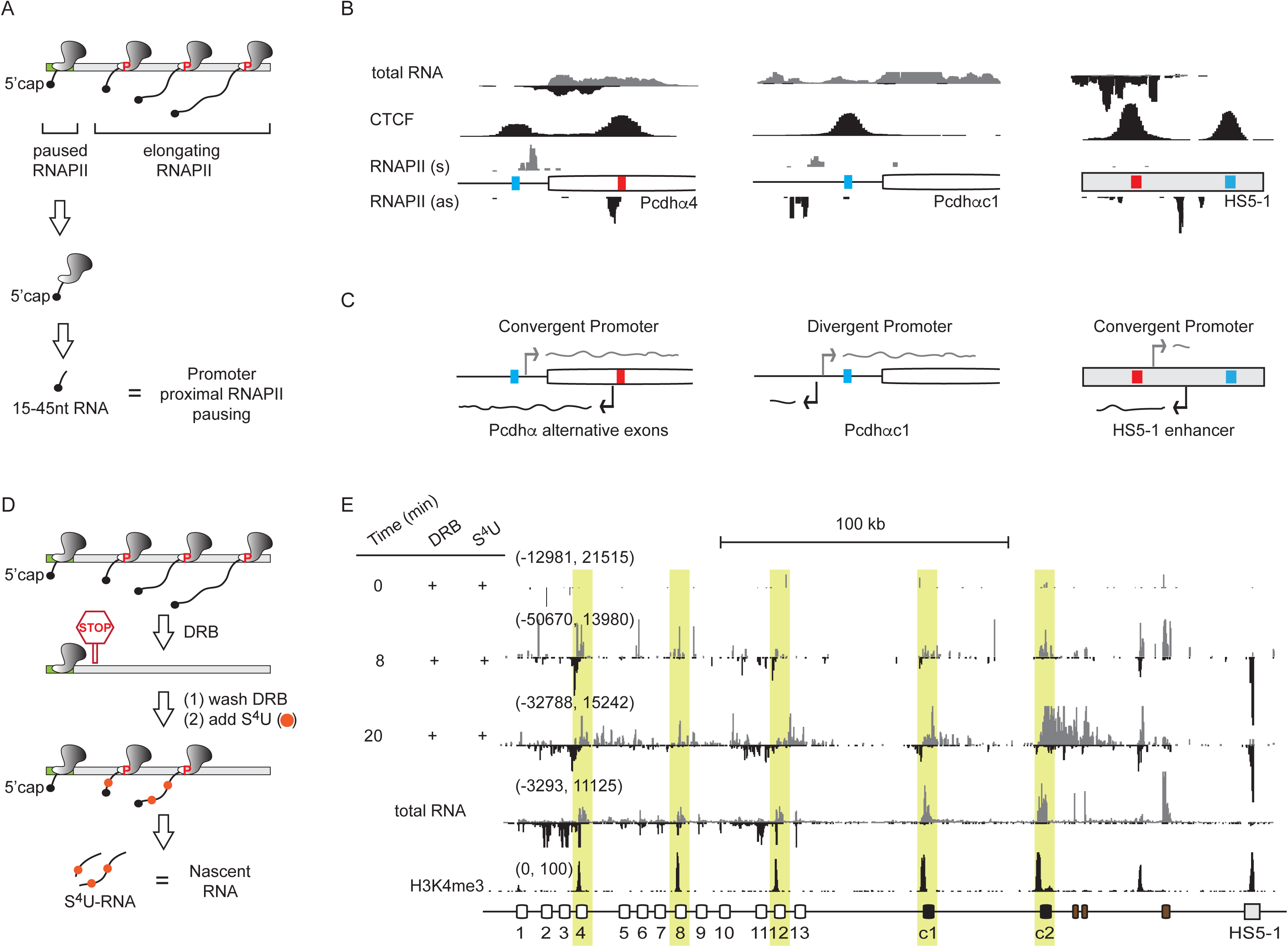
Convergent promoters in the Pcdhα alternative exons and HS5-1 enhancer. (A) Schematic diagram of Start-Seq. (B) Paused RNAPII relative to total RNA and CTCF binding.(C)Promoter architectures for Pcdhα4 (convergent), Pcdhαc1 (divergent) and the HS5-1 enhancer (convergent). (D) Schematic diagram of s^4^UDRB-cRNA-Seq. (E) Transcription by RNAPII relative to total RNA and H3K4me3 in SK-N-SH cells. The x-axis represents the linear sequence of the genomic organization of the Pcdhα cluster. The numbers on the left-hand side of each track represent the minimum and maximum densities in read per million.

Antisense convergent transcription is widespread in the mammalian genome. Yet, the activity of convergent transcribing polymerases remains unclear (see Discussion for more details). To assess RNAPII activity at the pCBS- and eCBS-promoters, we analyzed transcription in SK-N-SH cells using s^4^UDRB-Seq (Fuchs et al., 2014; Singh and Padgett, 2009) (Figure 2D). This method combines synchronization of RNAPII at promoters with incorporation of the nucleoside 4-thiouridine (s^4^U) during RNA synthesis. SK-N-SH cells were treated with 5,6-Dichloro-1-β-D-ribofuranosylbenzimidazole (DRB) to block RNAPII carboxy-terminal domain (CTD) phosphorylation, a step required to release paused RNAPII from promoters in their transition from initiation to productive elongation. DRB inhibition is reversible, and upon removal from the cell culture media, a wave of newly transcriptional elongating RNAPII leads to the incorporation of s^4^U into newly synthesized RNAs. s^4^U is rapidly incorporated into living cells without the need of cell lysis or nuclear isolation. Given the thiol-specific reactivity of s^4^U, s^4^U-labeled RNA can be covalently and reversibly captured and sequenced. Consistent with the Start-Seq data, we observed convergent, elongating RNAPII from both pCBS- and eCBS-promoters of α4, α8 and α12, and divergent RNAPII from the pCBS-proximal promoter of αc1 (Figure 2E and Figure S5). We also observed convergent elongating RNAPII at the HS5-1 enhancer, also consistent with the presence of convergent promoters (Figure 2E and Figure S5). Thus, there is a remarkable symmetry between the location of CTCF/Cohesin binding sites and sense and antisense transcription from the Pcdhα promoters and the HS5-1 enhancer. However, contrary to the sense and antisense RNA transcribed from a Pcdhα alternate exon, both enhancer RNAs are not polyadenylated and therefore appear to turn over faster (Figure S4 and S5).

Additional evidence for the presence of convergent promoters associated with both the pCBS and eCBS sequences was provided by the analysis of published ENCODE DNaseI hypersensitivity and ChIP-Seq data, which revealed the binding of distinct classes of transcription factors (TF) to the pCBS and eCBS sites of transcriptionally active alternate exons in SK-N-SH cells (Figure S6). Specifically, TFs belonging to the ETS family bind to the pCBS-promoter, while TFs belonging to the bHLH family bind to the eCBS-promoter (Figure S6). Interestingly, both these classes of TFs are implicated in regulating genes involved in neuronal development and differentiation, such as different members of cell-adhesion proteins (Hollenhorst et al., 2011).

### Transcription of antisense lncRNAs triggers activation of sense promoters

Understanding the architecture of the sense and antisense promoters of the Pcdhα genes and the HS5-1 enhancer provided the opportunity to address mechanistic questions regarding Pcdhα stochastic promoter choice. Specifically, we aimed to test the functional relationship between transcriptional activation of the sense and antisense promoters, and transcription of their respective RNAs. To do so, we designed a gain-of-function approach that allowed us to uncouple transcription of the antisense long noncoding RNA from transcription of the sense coding Pcdhα mRNA in the context of the endogenous Pcdhα gene cluster. Specifically, we made use of a catalytic-inactive CRISPR-dCas9 protein fused to a tripartite transcriptional activator (dCas9-VPR) (Chavez et al., 2015) to selectively activate the pCBS- or eCBS-promoters of silent Pcdhα genes (Figure 3A). We chose HEK293T cells, as in this cell line most Pcdhα genes, with the exception of Pcdh α10 and αc2, are silent. This made it possible to selectively activate Pcdh α4, α6, α9 and α12 (Figure S7). As expected, dCas9-VPR activation of the Pcdhα4 sense promoter resulted in robust synthesis of the Pcdhα4 s-cRNA (Figure 3B and 3C). By contrast, activation of the Pcdhα4 antisense promoter resulted not only in high levels of transcription of antisense RNA, but high levels of sense RNA was also observed (Figure 3B and 3C). This pattern of sense and antisense RNA transcription observed upon eCBS-promoter activation mirrored that observed in primary neurons and SK-N-SH cells at steady-state (Figure 1D and Figure S4). Similar to Pcdhα4 gene, activation of the eCBS-promoters of the Pcdh α6, α9, and α12 also resulted in transcription of both sense and antisense RNAs (Figure 3D).

**Figure 3:**
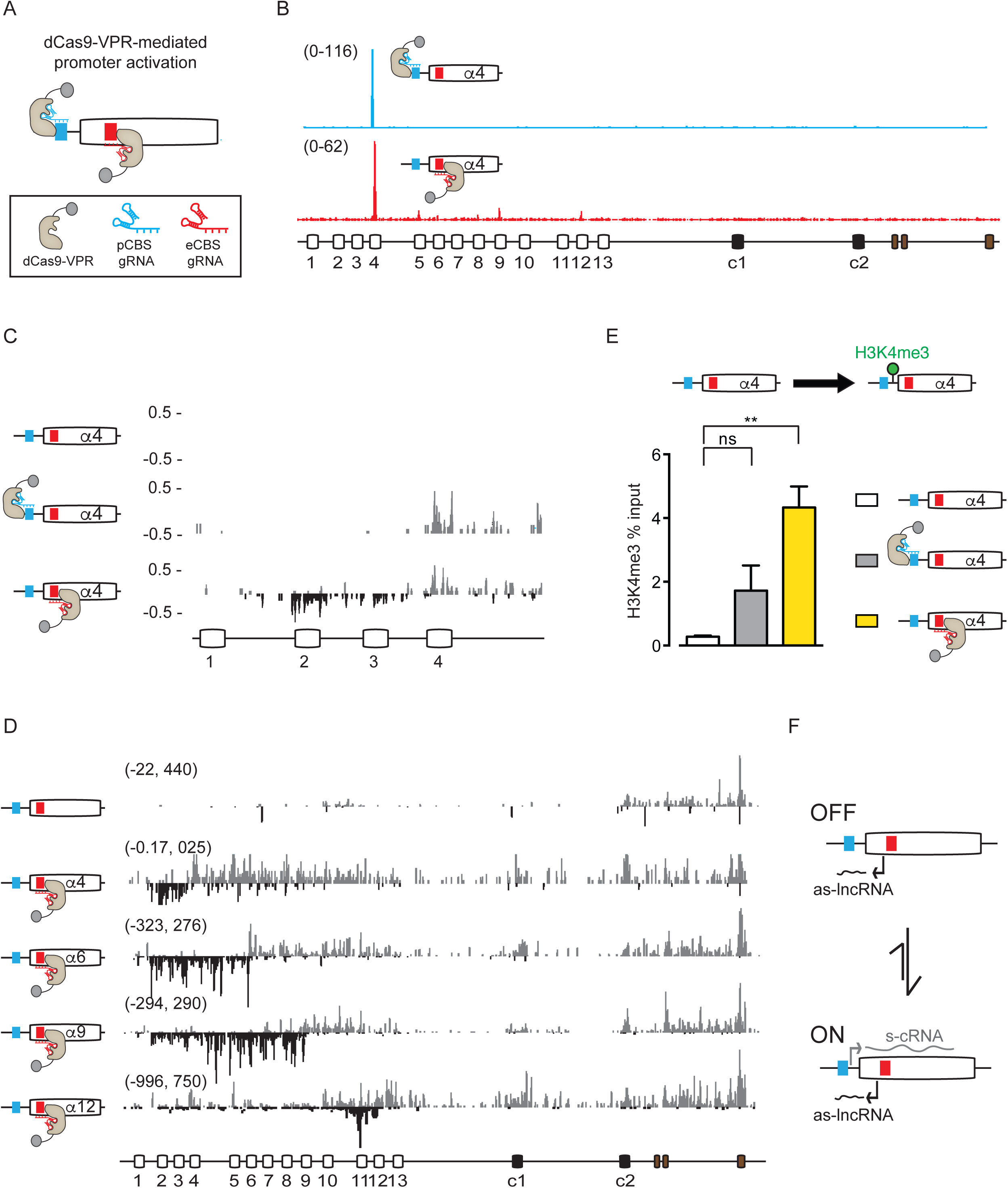
Transcription of the antisense lncRNA triggers activation of sense promoters. (A) Schematics of dCas9-VPR-mediated activation of pCBS- and eCBS-promoters. (B) Distribution of dCas9-VPR targeted to the pCBS-promoter (blue) and the eCBS-promoter (red) of Pcdhα4 in HEK293T cells assayed by ChIP-Seq. (C) Activation of the sense and antisense promoters in Pcdhα4 by dCas9-VPR. (D) Transcription of sense and antisense RNA upon activation of the eCBS-promoters of Pcdh α4, α6, α9, α12 by dCas9-VPR. The data for Pcdh α6, α9, α12 and no activation are cRNA-Seq. The data for Pcdh α4 are RNA-Seq. (E) Enrichment of H3K4me3 at the Pcdhα4 promoter measured by ChIP-qPCR. (F) Schematic diagram for how transcription of the as-lncRNA from an eCBS-promoter leads to activation of pCBS sense promoter and transcription of the s-cRNA. For (B-D), the x-axis represents the linear sequence of the genomic organization of the Pcdhα cluster. For (B) and (D), the numbers on the left-hand side of each track represent the minimum and maximum densities in read per million. Those numbers are indicated as the y-axis. For (E), errors (n=3) represent s.e.m. and statistical significance was calculated with a Student unpaired *t*-test.

These observations suggest the possibility that transcription of the antisense RNA by the eCBS-promoter activates its cognate pCBS-promoter to allow for transcription of the sense coding RNA. To test this model, we measured the levels of histone H3 lysine 4 trimethylation (H3K4me3) at the Pcdhα4 sense promoter. This histone post-translational modification is a marker for transcriptionally active promoters and is enriched between two CBS sites of Pcdhα active exons in SK-N-SH cells (Figure 1D). We therefore carried out chromatin immunoprecipitation studies followed by quantitative PCR (ChIP-qPCR) and observed an increase in H3K4me3 at the Pcdhα4 sense promoter upon transcription the antisense RNA by dCas9-VPR (Figure 3E).

Taken together, these data are consistent with a model in which transcription of the antisense RNA by the eCBS-promoter activates its cognate pCBS-promoter, thus generating convergent sense and antisense transcripts (Figure 3F). This level of exquisite specificity is remarkable, considering that the as-lncRNA transcribes through multiple upstream pCBS-promoters (Figure 1D and 3D), yet the only pCBS-promoter that is activated is the one proximal to the site of initiation of the antisense RNA. We speculate that this proximal specificity is a consequence of functional coupling between transcription and RNA processing mediated by the carboxy-terminal (CTD) of the RNAPII and the cap-binding complex (Maniatis and Reed, 2002).

### Antisense transcription leads to CTCF binding and promoter/enhancer long-range DNA interactions

The expression of Pcdhα sense transcripts correlates with the binding of CTCF and Cohesin proteins to the pCBS and eCBS sites and to long-range DNA looping between active promoters and the HS5-1 enhancer (Guo et al., 2012; 2015). Using ChIP-Seq, we observed that both CBSs of the inactive Pcdh α4, α6, α9, and α12 in the HEK293T used in this study are not occupied by CTCF (Figure 4A). We therefore asked whether antisense transcription promotes the binding of the CTCF protein to its sites on the activated exon. To accomplish this, we again turned to the dCas9-VPR-based gain-of-function strategy described above and illustrated in Figure 3A. Consistent with the mechanistic coupling of promoter activation and CTCF/Cohesin binding (Guo et al., 2012; Monahan et al., 2012), we observed a statistically significant enrichment of CTCF occupancy at both pCBS and eCBS sites upon dCas9-VPR activation of their antisense promoters relative to the activation of their sense promoters (Figure 4B).

**Figure 4:**
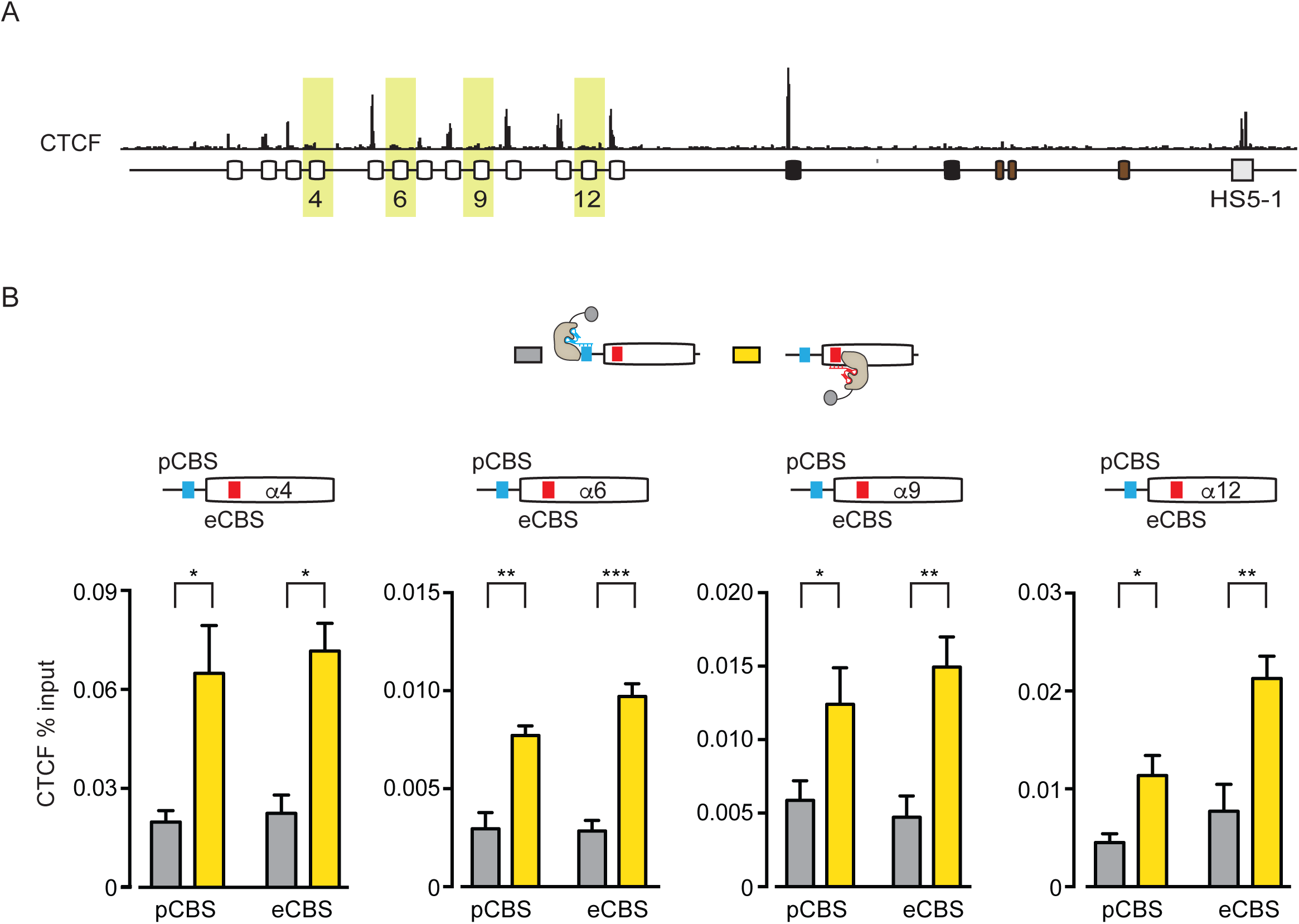
Antisense lncRNA transcription leads to CTCF binding. (A) Distribution of CTCF in HEK293T cells assayed by ChIP-Seq. The exons free of CTCF and activated in this study are highlighted in yellow. (B) Enrichment of CTCF occupancy at the pCBS and the eCBS sites of Pcdh α4, α6, α9, α12 assayed by ChIP-qPCR. Errors (n>3) represent s.e.m. and statistical significance was calculated with a Student unpaired *t*-test.

The binding of CTCF suggested the possibility that antisense transcription from the activated exon leads to CTCF/Cohesin-dependent long-range DNA looping between the active promoter and the HS5-1 enhancer. To address this possibility, we focused on the Pcdhα12 exon. We performed triplicate *in situ* cHi-C experiments on HEK293T cells transfected with dCas9-VPR to activate either the Pcdhα12 pCBS- or eCBS-promoter and obtained a combined contact matrices at 2kb resolution (Figure 5A). Analysis of *in situ* cHi-C data from Pcdhα12 pCBS-activated HEK293T cells showed three long-range DNA contacts corresponding to the interactions between Pcdh α10, α11 and α13 and the HS5-1 enhancer (Figure 5B, left). All three of these exons are indeed stably bound by CTCF in our HEK293T cells (Figure 4A). Most importantly, we did not observe a long-range DNA interaction between the Pcdhα12 and the HS5-1 enhancer (Figure 5B, left). By contrast, analysis of *in situ* cHi-C data from Pcdhα12 eCBS-activated HEK293T cells revealed the formation of newly well-defined long-range DNA interactions between the activated Pcdhα12 promoter/exon and the HS5-1 enhancer, and a concomitant reduction of contacts between the Pcdh α10, α11 and α13 and the HS5-1 enhancer (Figure 5B, middle). Consistent with the formation of a Pcdha12/HS51 complex, we observed an increase of CTCF binding at the HS5-1 enhancer in HEK293T cells activated with dCas9-VPR targeting the eCBS-compared to the pCBS-promoter (Figure S8). Most importantly, the same type of promoter/enhancer landscape observed here is characteristic of that observed for the SK-N-SH cells (Figure S3C and Figure 5B, right). As described above, in these cells, Pcdhα12 expression is epigenetically stable and both its pCBS and eCBS sites are bound by the CTCF protein and the Cohesin complex (Figure 1D) and in contact with their corresponding L- and R-CBSs on the HS5-1 enhancer (Figure S3C).

**Figure 5:**
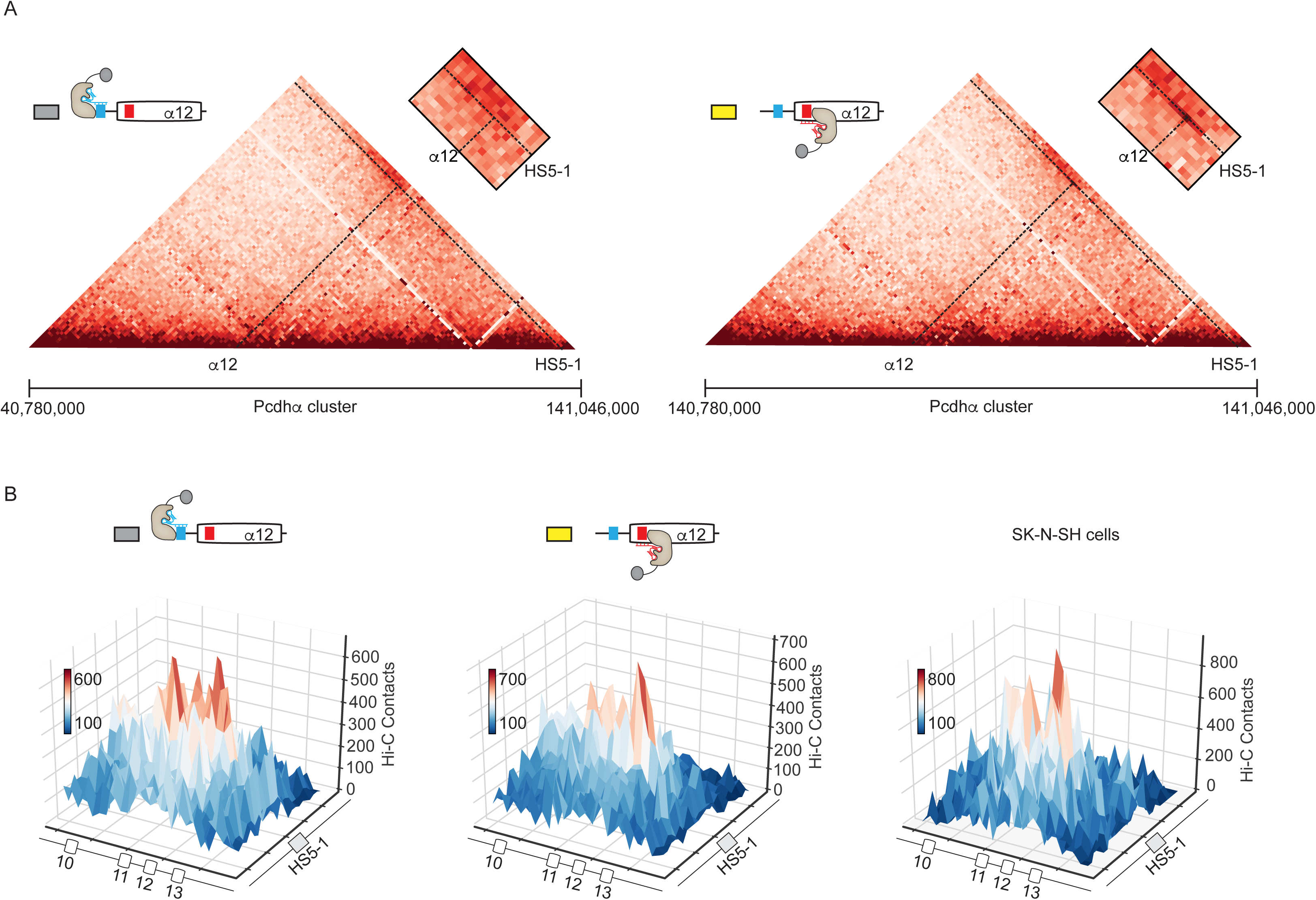
Antisense lncRNA transcription leads to promoter/enhancer DNA interactions. (A) *In situ* cHi-C contact maps at 2kb resolution for the Pcdhα cluster in HEK293T cells activated with dCas9-VPR targeting the pCBS-promoter (Left) or the eCBS promoter (Right) of Pcdhα12. Coordinates: chr5:140,780,000-141,046,000, hg38. The position of the Pcdhα12 exon and the HS5-1 enhancer are indicated by the dotted lines. The inset is a zoom-in of the region surrounding the Pcdhα12 exon in complex with the HS5-1 enhancer. (B) Local peak analysis of *in situ* cHi-C long-range interactions between Pcdhα promoters and the HS5-1 enhancer. *In situ* cHi-C of HEK293T cells activated with dCas9-VPR targeting the pCBS promoter (Left) or the eCBS promoter (Middle) of Pcdhα12. *In situ* cHi-C of SK-N-SH cells (Right).

Together, these data suggest that binding of the CTCF protein and formation of long-range DNA looping between active Pcdhα promoters and the HS5-1 enhancer are coupled and regulated by a loop extrusion mechanism. We propose that, as previously described for other promoter-enhancer complexes in the mammalian genome (Fudenberg et al., 2016), long-range DNA looping between the active promoters and the HS5-1 enhancer occurs as a consequence of stalled Cohesin complexes by extrusion barriers. According to this model, CTCF bound to DNA acts as such an extrusion barrier, and therefore stalls actively translocating Cohesin molecules at the activated Pcdhα promoters. By contrast, the inactive Pcdhα promoter, which is not bound by CTCF, is not brought into contact with the HS5-1 enhancer.

### Antisense lncRNA transcription leads to the conversion of 5mC to 5hmC at Pcdhα promoters

The data presented thus far suggest a model in which the stochastic choice of Pcdhα alternate promoters requires coupling between transcription of antisense lncRNAs and the assembly of a promoter/enhancer complex by CTCF/Cohesin. However, it is not clear how transcription of the antisense lncRNA leads to the recruitment of the CTCF/Cohesin complex, the assembly of a functional promoter/enhancer complex, and stable transcriptional activation of a Pcdhα sense coding RNA. As mentioned above, a key difference between Pcdhα active and inactive sense promoters is that the former display low levels of CpG methylation by the pCBS and eCBS sites, while the later are highly methylated (Guo et al., 2012; Kawaguchi et al., 2008; Tasic et al., 2002). Given this correlation, and the observation that DNA methylation of the CTCF binding sites blocks CTCF binding (Bell and Felsenfeld, 2000), a potential mechanism of CTCF/Cohesin binding to exons following as-lncRNA transcription is demethylation of the methylated CpG sequences.

In mammals, 5-methylcytosine (5mC) modified CpG sequences are converted to unmodified cytosine (C) by the activity of TET deoxygenase enzymes, which mediate the oxidation of 5mC to 5-hydroxymethylcytosine (5hmC), 5-formylcytosine (5fC) and 5-carboxylcytosine (5caC) (Wu and Zhang, 2017). Thymine DNA glycosylase (TDG) then converts 5caC to C by a base excision repair mechanism (Wu and Zhang, 2017). To test the possibility that transcription of the as-lncRNA leads to demethylation of CpG islands, we measured the levels of 5mC and 5hmC for the Pcdhα4 and the Pcdhα12 promoter/exons in HEK293T cells by Methylated DNA Immunoprecipitation (MedIP) upon dCas9-VPR mediated activation of their respective sense (pCBS) and antisense (eCBS) promoters. Consistent with our hypothesis, activation of the Pcdh α4 and α12 eCBS-promoters resulted in a decrease of 5mC/5mhC levels at the pCBS and the eCBS sites (Figure 6A and 6B). By contrast, activation of the Pcdhα4 and the Pcdhα12 pCBS-promoter resulted in only a statistically significant decrease of 5mC/5mhC levels by the pCBS but not the eCBS (Figure 6A and 6B). These data are consistent with the hypothesis that transcription of the antisense lncRNA promotes CpG DNA conversion from 5mC to 5hmC of both pCBS and eCBS sites, and suggest that demethylation of both sites is a required step for the stable binding of CTCF to its DNA binding sites.

**Figure 6:**
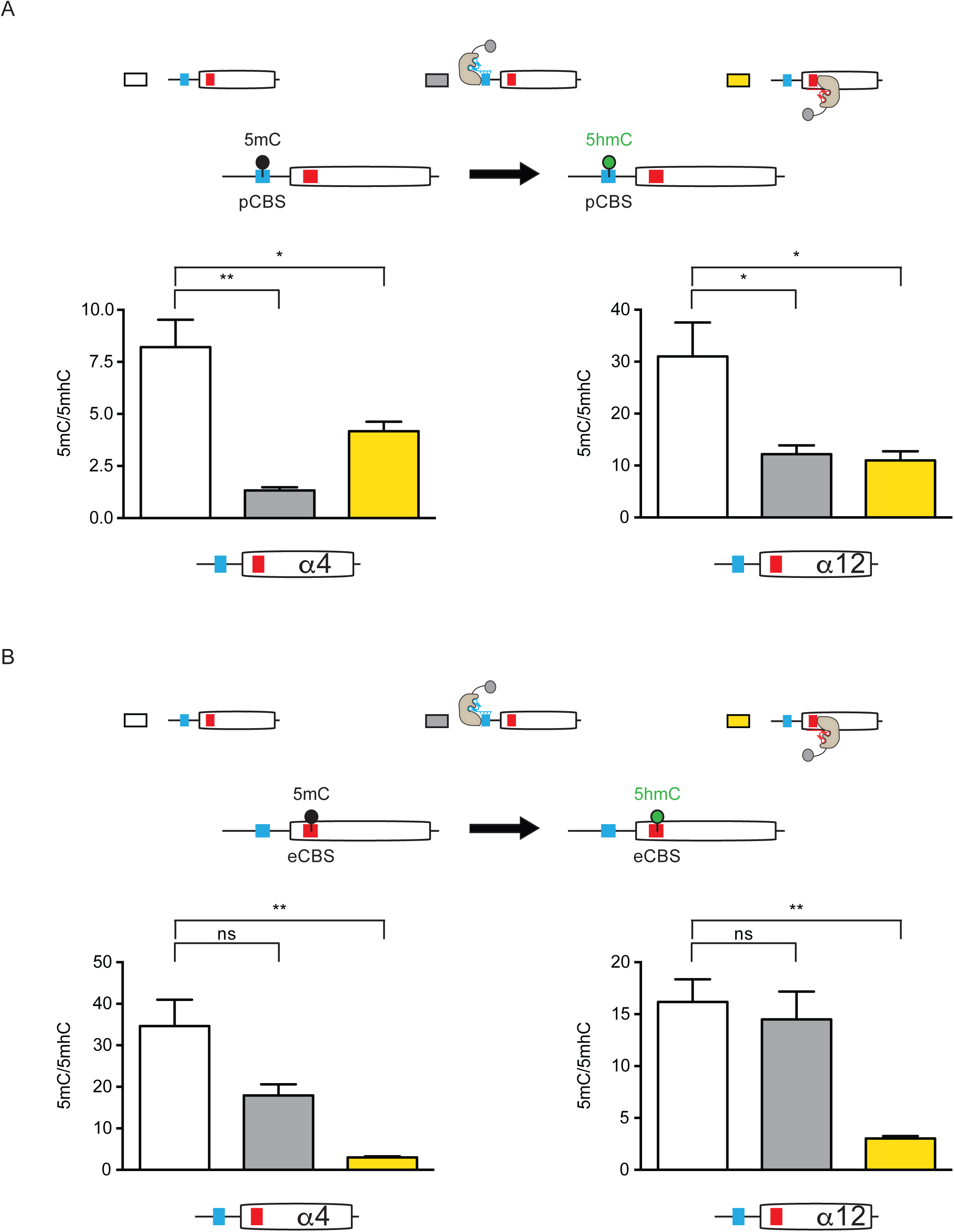
Antisense lncRNA transcription leads to the conversion of 5mC to 5hmC. (A) Relative levels of 5mC and 5hmC at the pCBS of Pcdhα4 and Pcdhα12. (B) Relative levels of 5mC and 5hmC at the eCBS of Pcdhα4 and Pcdhα12. Levels of 5mC and 5hmC are assayed by MeDIP-qPCR. Errors (n=3) represent s.e.m. and statistical significance was calculated with a Student unpaired *t*-test.

### Conversion of 5mC to 5hmC at Pcdhα promoters correlates with their activation *in vivo*

The data presented above are consistent with a model in which the ground state of a Pcdhα promoter DNA is methylated, and conversion of 5mC to 5hmC, targeted by transcription of an antisense lncRNA, controls Pcdhα sense promoter activation. To test this model *in vivo*, we made use of the mouse main olfactory sensory epithelium (mOE), as an *in vivo* developmental system to study the relationship between promoter DNA methylation and Pcdhα gene expression. Previous studies have shown that the Pcdh gene cluster is stochastically and combinatorially expressed in OSNs, and that the Pcdhα genes play a fundamental role in OSN wiring (Hasegawa et al., 2008; 2016; Mountoufaris et al., 2017). We re-analyzed recent published work carried out to characterize the levels of 5mC and 5hmC in the three cell types that represent discrete neurodevelopmental stages in the mOE: horizontal basal cells (iCam1^+^), immediate neural precursors (Ngn1^+^) and mature olfactory sensory neurons (Omp^+^) (Colquitt et al., 2013). Horizontal basal cells are quiescent multipotent cells that produce all of the cell types present in the mOE. Immediate neural precursors are post-mitotic cells precursors to olfactory sensory neurons. Olfactory sensory neurons are terminally differentiated primary sensory neurons. Consistent with our model, we found that clustered Pcdhα alternate exons and their promoters are enriched in 5mC in iCam1^+^ cells indicating that the ground state of all Pcdhα alternate promoters is methylated and repressed (Figure 7A and 7B). However, with the development of olfactory sensory neurons (iCam1^+^ → Ngn1^+^ → Omp^+^), we observed an increase of 5hmC in the Pcdhα alternate promoters and exons (Figure 7A and 7B). To test whether conversion of 5mC to 5hmC is accompanied by activation of Pcdhα promoter and their expression, we performed RNA-Seq experiments in iCam1^+^, Ngn1^+^ and Omp^+^ cells. Consistent with our hypothesis, conversion of 5mC to 5hmC correlated with the expression of both antisense long noncoding and sense coding Pcdhα RNAs (Figure 7C). Finally we tested whether Pcdhα expression is accompanied by the formation of long-range DNA contacts between the Pcdhα promoters and the HS5-1 enhancer *in vivo*, and performed *in situ* Hi-C experiments in iCam1^+^, Ngn1^+^ and Omp^+^ cells. Consistent with our model, we observed a remarkable increase in alternate promoters/HS5-1 enhancer interactions during neuronal differentiation of the mOE (Figure 7D and Figure S9).

**Figure 7:**
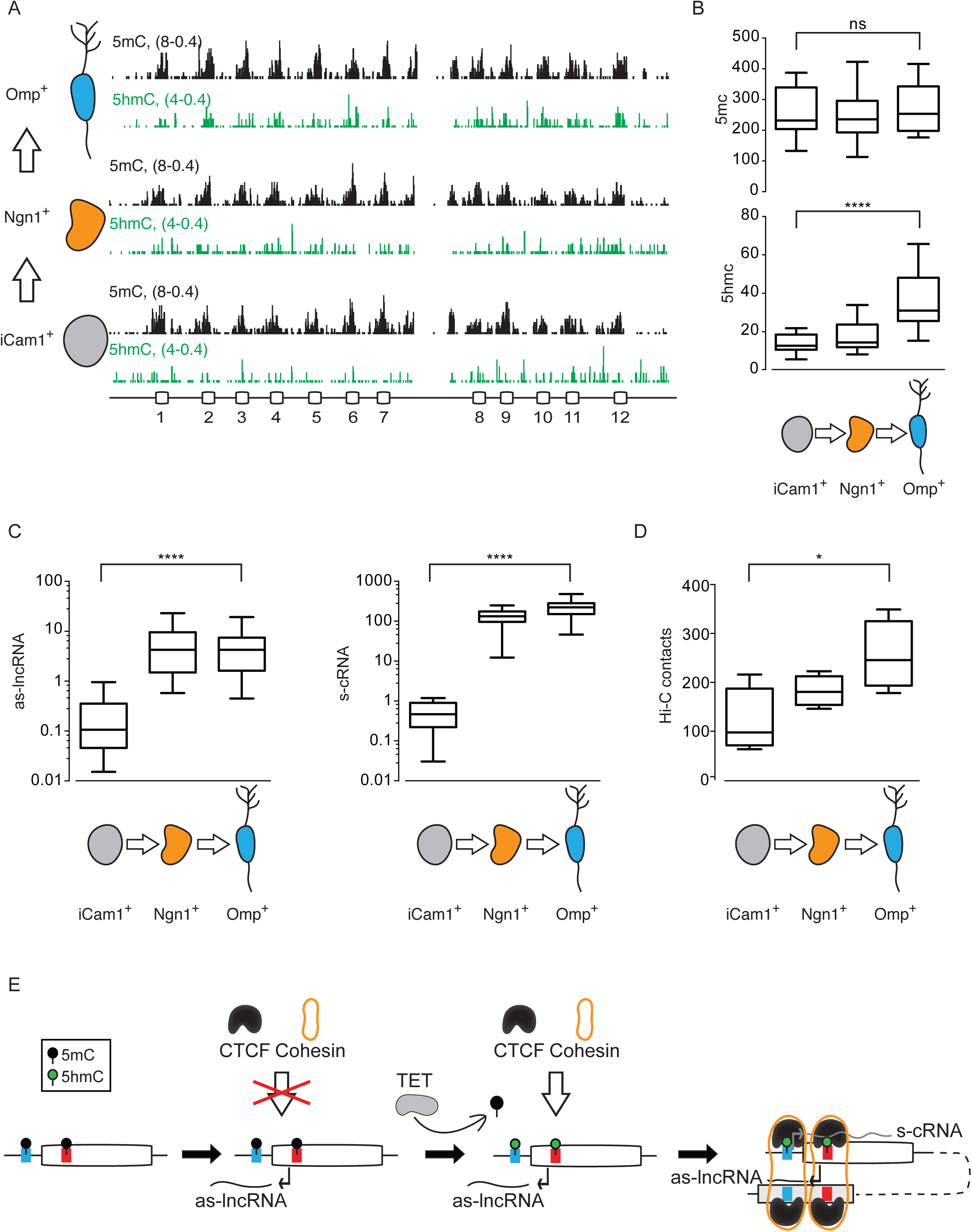
Conversion of 5mC to 5hmC at Pcdhα promoters correlates with their activation *in vivo*. (A) 5mC (Black) and 5hmC (Green) profiles of the Pcdhα alternate promoters and exons in horizontal basal cells (iCam1^+^), immediate neural precursors (Ngn1^+^) and mature olfactory sensory neurons (Omp^+^) of the mouse main olfactory epithelium. The x-axis represents the linear sequence of the genomic organization of the mouse Pcdhα cluster. The numbers on the left-hand side of each track represent the minimum and maximum read densities in read per million. (B) Average of cumulative RPM values for the Pcdhα alternate promoters/exons for 5mC (top) and 5hmC (bottom). (C) Average of cumulative RPM values for the Pcdhα antisense lncRNA (as-lncRNA) and sense coding RNA (s-cRNA). (D) Average of cumulative *in situ* Hi-C contacts for the Pcdhα alternate promoters/exons. (E) Model of which transcription of a lncRNA triggers activation of Pcdhα promoter and expression of its mRNA by mediating DNA demethylation of the pCBS and eCBS, a step required for the binding of the CTCF/Cohesin protein complex. CTCF/Cohesin binding promotes the assembly of a promoter/HS5-1 enhancer complex. For (B-D), data are represented in Box and whiskers. Error bars represent minimum and maximal values and statistical significance was calculated with one-way ANOVA.

## DISCUSSION

Stochastic, combinatorial expression of individual Pcdh protein isoforms in purkinje (Esumi et al., 2005) and olfactory sensory neurons (Mountoufaris et al., 2017), as revealed by single-cell RNA expression studies, generates distinct combinations of Protocadherin isoforms that function as a cell-surface identity code for individual neurons. This code is required for self-avoidance of sister neurites, a process critical for normal neuronal connectivity and neurite patterning (Lefebvre et al., 2012; Mountoufaris et al., 2017). The extraordinary functional diversity of Pcdhs required for self-avoidance is a consequence of a mechanism of stochastic promoter activation of alternate exon promoters, which occurs independently in each of the three gene clusters, and on both allelic chromosomes. This diversity is further expanded through nearly random dimerization of distinct Pcdh isoforms, the unique anti-parallel interactions of homophilic Pcdh complexes at the cell surface, and a stringent homophilic specificity (Rubinstein et al., 2017). In contrast to purkinje and olfactory neurons, however, serotonergic neurons require only a single Pcdh identity provided by the expression of the c-type, Pcdhαc2 protein, which functions in axonal tiling (Chen et al., 2017; Katori et al., 2017). Thus, the establishment of differential transcriptional programs regulating the expression of the alternate and c-type Pcdh gene clusters isoforms enables distinct Pcdh functions in different neuronal cell types. Uncovering such transcriptional programs will reveal fundamental principles of neural circuit assembly and the role of Pcdh-mediated neural cell-surface diversity in the mammalian brain. Here we identify a mechanism by which Pcdhα alternate exon promoters are stochastically activated in individual neurons, and discuss the possibility that this mechanism may function more broadly in promoter choice and gene expression in vertebrates.

### Insights into the mechanism of stochastic promoter choice

Here, we provide evidence that stochastic activation of individual Pcdhα alternate promoters requires mechanistic coupling between transcription of an antisense lncRNA and DNA looping between individual Pcdhα promoters and the HS5-1 enhancer (Figure 7E). This mechanism requires the binding of the CTCF/Cohesin complex to both the promoter and enhancer. Specifically, each Pcdhα alternate exon is characterized by convergent promoters in close proximity to the two CTCF/Cohesin binding sites (pCBS and eCBS). The eCBS-promoter initiates transcription of a long noncoding RNA that extends through the pCBS-promoter and into upstream intergenic sequences, leading to transcriptional activation of the pCBS-promoter. This process is accompanied by the conversion of 5-methylcytosine (5mC) to 5-hydroxymethylcytosine (5hmC) of the pCBS and eCBS sequences. While 5mC is usually associated with transcriptionally inactive genes, 5hmc is associated with gene activation (Colquitt et al., 2013; Ficz et al., 2011; Kriaucionis and Heintz, 2009). This epigenetic switch from 5mC to 5hmC allows for the binding of the CTCF protein to its dual binding sites. CTCF, together with the Cohesin complex, mediates long-range DNA interactions between the active promoter and the Pcdhα cluster-specific enhancer, HS5-1. Formation of a promoter/enhancer complex commits the Pcdhα promoter choice and thus facilitates transcription of a Pcdhα mRNA.

A fundamental question raised by these observations is how antisense promoters are stochastically activated in individual neurons during development. Given the observation that the ground state of the Pcdhα gene cluster is inactive and marked by 5mC (Figure 7A and 7B), we speculate that activation of eCBS-promoters in the Pcdhα gene cluster is regulated by the activity of transcription factors capable of binding methylated DNA. In contrast to the Pcdhα gene cluster, the alternate exons in the Pcdh β and γ clusters bear a single CBS site in their promoters (pCBS), but lack a CTCF/Cohesin binding site and the antisense promoter in the downstream exon and therefore do not transcribe antisense lncRNAs. Nevertheless, Pcdh β and γ promoter choice is stochastic (Esumi et al., 2005; Mountoufaris et al., 2017) and transcriptional enhancer elements, similar to the HS5-1 enhancer, located distal to the Pcdhγ gene cluster have been proposed to be required for their transcription (Yokota et al., 2011). Thus, the mechanism of random promoter choice in these gene clusters remains unknown, and is likely to be cell type-specific. Indeed, by contrast to the Pcdhα gene cluster, which is expressed exclusively in the nervous system, the Pcdh β and γ gene clusters are expressed more broadly (https://www.gtexportal.org/home/).

### The molecular logic of convergent promoters

Bi-directional RNA transcription is a common feature of mammalian promoters and enhancers (Wu and Sharp, 2013; Core et al., 2014; Henriques, et al, 2018). The transcripts can be divergent, thus non-overlapping, or, as is the case of the Pcdhα gene cluster, convergent, which produces overlapping complementary RNAs. Divergent transcription at promoters usually produces upstream non-coding RNAs, transcribed toward the 5’ end of the gene, that are on average 50-2000 nucleotides long and relatively unstable (Wu and Sharp, 2013). Divergent transcription is widespread in mammals and is characteristic of active promoters that lack a distinct TATA box motif and are enriched of CpG islands (Wu and Sharp, 2013). By contrast, convergent transcription can produce long and stable antisense noncoding RNAs that often overlap with the sense coding RNA (Brown et al., 2018; Mayer et al., 2015). These antisense RNAs can function to either activate or repress transcription of the coding RNA from the sense promoter, in a process known as transcription interference (Bonasio and Shiekhattar, 2014). Interestingly, contrary to divergent promoters, active genes marked by convergent transcription are characterized by a unique chromatin signature (Brown et al., 2018; Mayer et al., 2015; Murray and Mellor, 2016; Scruggs et al., 2015). In this case, antisense RNA transcription has been proposed to actively shape a unique chromatin environment necessary to promote transcription of the cognate sense RNA. The example of convergent transcription described here suggests a model in which noncoding antisense RNA transcription plays a fundamental role in stochastic promoter activation by establishing an epigenetically stable chromatin architecture at Pcdhα promoters, which is critical to the formation of proper promoter/enhancer long-range DNA looping. This mechanism involves coupling between RNAPII, a DNA deoxygenase TET enzyme and the insulator complex CTCF/Cohesin. We note that a transcription-dependent mechanism of transcriptional activation is not unprecedented. Specifically, transcription of the tumor suppressor gene, TCF21, was shown to be activated by an antisense RNA whose transcription is initiated at an intronic promoter sequence located within the TCF21 gene (Arab et al., 2014). Like the mechanism proposed here, transcription through the TCF21 promoter leads to TET-mediated DNA demethylation and activation of the TCF21 sense strand promoter. Here, we show that this mechanism is used for stochastic choice of Pcdhα promoters, which has profound implications regarding neuronal circuit assembly during development.

### De-repression followed by stochastic enhancer-promoter interactions as a general mechanism for transcriptional activation

Since we used the differentiating mouse olfactory epithelium as an *in vivo* model system for stochastic Pcdhα gene activation, we could not ignore the striking similarities in the regulatory logic between Pcdhα and olfactory receptor (OR) promoter choice. In both cases, the ground state of the stochastically chosen promoters is repressed and inaccessible to transcriptional activator proteins. In the case of the Pcdhα gene cluster, this repression is mediated predominantly by DNA methylation (Tasic et al., 2002; Toyoda et al., 2014), while OR genes are repressed by the assembly of constitutive heterochromatin (Magklara et al., 2011; Monahan et al., 2017). In both of these cases, however, repressive DNA or histone modifications are replaced by activating marks, concomitantly with selective binding of transcription factors that recruit distant transcriptional enhancers to the transcriptional start site. Because all the Pcdhα genes are clustered in a single chromosome, stochastic Pcdhα choice can be accomplished with cis-acting enhancers only. This is in contrast to OR gene choice, which appears to require the formation of a multi-chromosomal multi-enhancer hub that activates only one out the 2800 OR alleles distributed throughout the genome (Horta et al., 2018; Markenscoff-Papadimitriou et al., 2014; Monahan et al., 2018). Most likely, reliance on *cis* versus *trans* interactions also explains why the two systems require different molecular players to achieve transcriptional stochasticity: in the case of Pcdhα genes, CTCF and Cohesin are ideal to promote stochastic enhancer/promoter interactions since the loop extrusion mechanism will essentially allow the HS5-1 enhancer to scan the genome locally for the most proximal promoter bound by CTCF. In contrast, OR enhancers cannot deploy loop extrusion mechanisms to activate OR transcription because this process cannot accommodate *trans* interactions, explaining the absence of CTCF and Cohesin from OR enhancers and promoters (Monahan et al., 2018). Consequently, as Pcdhα choice relies on stable CTCF promoter binding, DNA methylation/hydroxymethylation provides an effective mechanism for stochastic promoter activation. An important consequence of this mechanism is that, since antisense transcription and hydroxymethylation appear to occur in a stochastic fashion, and independently of the HS5-1 enhancer, loop extrusion will not create a bias toward the selection of the Pcdhα promoter most proximal to the enhancer (Pcdhα13 and Pcdhα12 in human and mouse, respectively). Instead, loop extrusion will identify the promoter that is bound by CTCF, providing an elegant mechanism to overcome selection biases driven by genomic proximity. Therefore, it would not be surprising if other clustered genomic systems use the same principle to assure that distant DNA elements are utilized as frequently as proximal ones, with prime example the process of V(D)J recombination, whereby proximity bias would be detrimental to the generation of the most diverse immunoglobulins and T cell receptors possible.

## AUTHORS CONTRIBUTIONS

D.C. and T.M. identified, developed and addressed the core questions regarding promoter choice. D.C. performed the bulk of the experiments with substantial help from C.L.N and help from R.M.S. and E.L.C. A.H. and S.L. helped to develop the chromosome conformation studies, and A.H. performed the experiment with the help of D.C. and C.L.N and analyzed the data. E.E.D. and M.D.S. helped develop the RNAPII elongation studies and E.E.D. and D.C. performed the experiments. R. D. helped D.C. performing RNA sequencing experiments in the mouse olfactory epithelium. D.C. and T.M. wrote the paper with the help of all the authors.

## ACKNOWLEDGEMENTS

We thank Ira Schieren for assistance with flow cytometry. We thank Dr. Axel, Dr. Zuker, Dr. Max Gottesman. David Hirsh, Dr. Germano Cecere, Dr. Kevin Monahan, Dr. Mountoufaris and members of the Maniatis, Lomvardas and Simon labs for helpful discussions and suggestions on the manuscript. D.C. would like to thank Sean O'Keeffe for his training in bioinformatics analysis of the data and Dr. Karen Adelman for advice with the Start-Seq experiments. D.C. and T.M. would like to thank Dr. Ye Zhang and Dr. Ben Barres for the generous gift of human brain neurons and Dr. Victor Lobanenkov and Dr. Elena Pugacheva for the generous gift of the CTCF monoclonal antibody. This work was supported by the Helen Hay Whitney Postdoctoral Fellowship and NIH Path to Independence Award K99/R00 K99GM121815 (D.C.), and the NIH grants 1R01MH108579 and 5R01NS088476 (T.M.).

## SUPPLEMENTAL FIGURE LEGENDS

**Figure S1:**
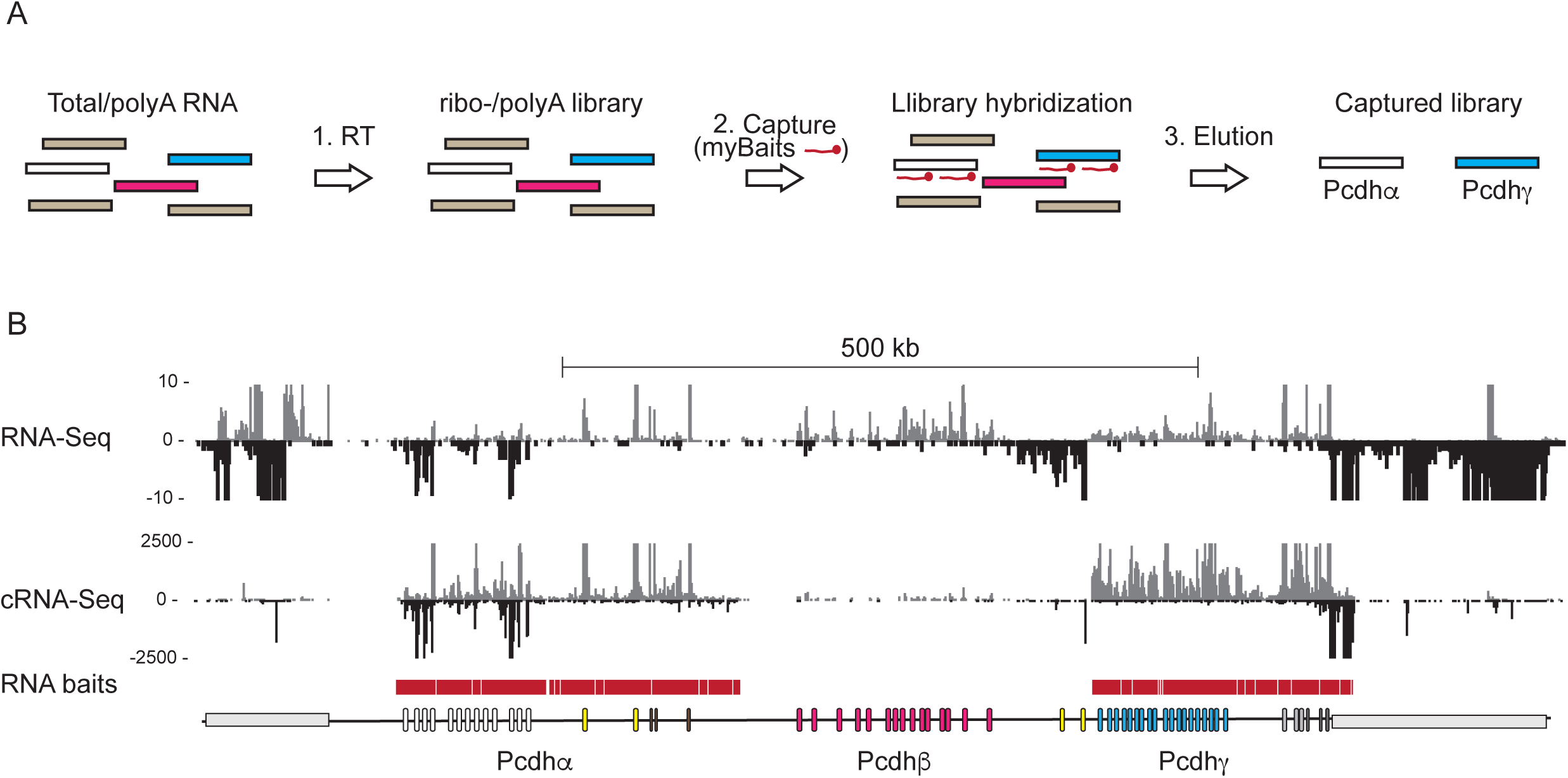
RNA-Sequencing and Capture RNA-Sequencing. (A) Schematic diagram of Capture RNA-Sequencing (cRNA-Seq). The white, pink and blue bars indicate RNA from the Pcdh α, β and γ clusters, respectively. The brown bars indicates RNA from the rest of the genome. (B) RNA-Seq and cRNA-Seq from total RNA isolated from SK-N-SH cells. Red bar: myBaits for the Pcdh α and γ clusters.

**Figure S2:**
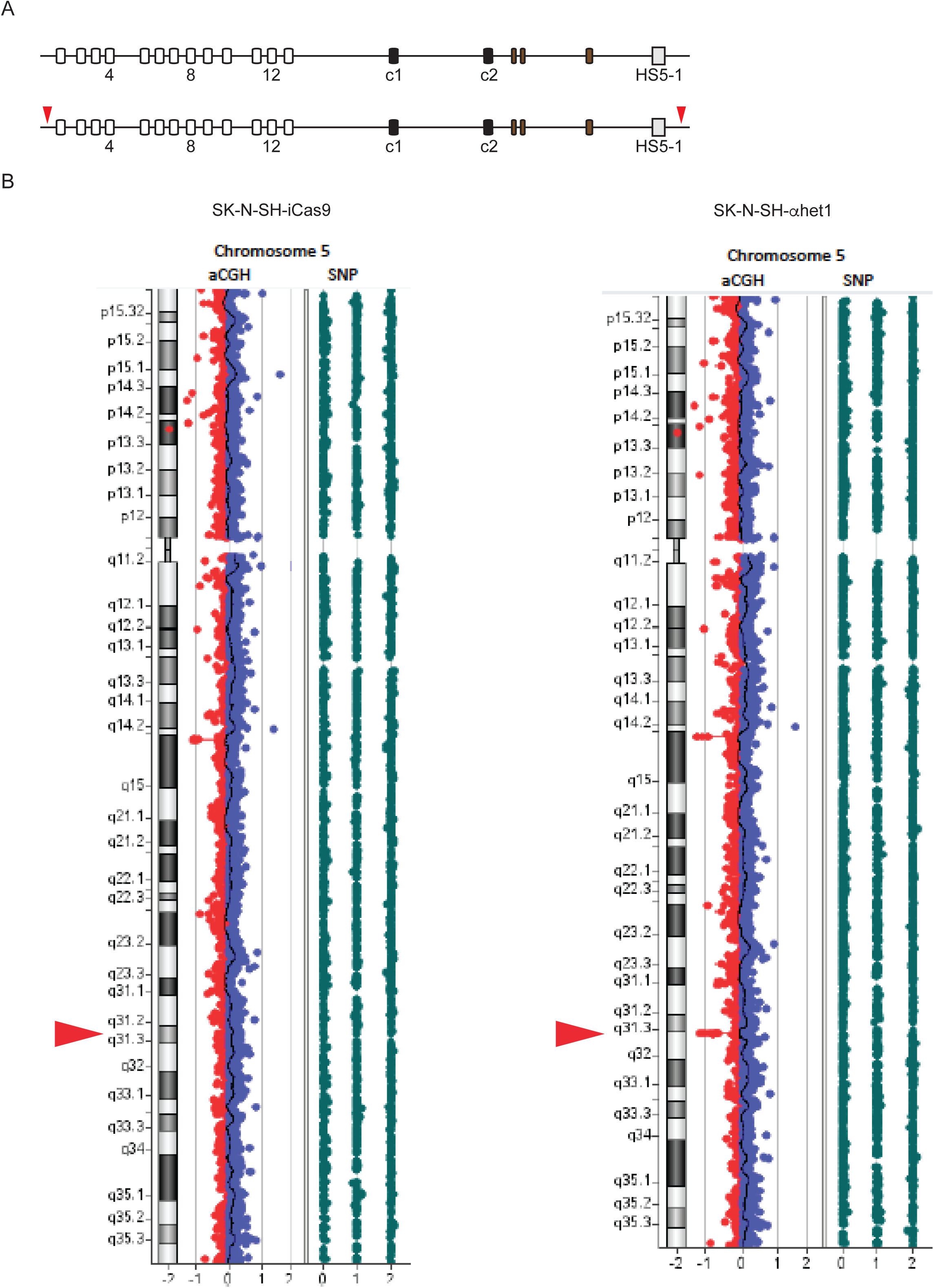
Generation of a SK-N-SH cell heterozygous for the Pcdhα gene cluster. (A) Pcdhα cluster. Red arrows indicate the location of the gRNAs used to delete a copy of the Pcdhα cluster. (B) Array Comparative Genomic Hybridization (aCGH) analysis confirms the deletion of a copy of a Pcdhα cluster in the SK-N-SH-αhet1 cells.

**Figure S3:**
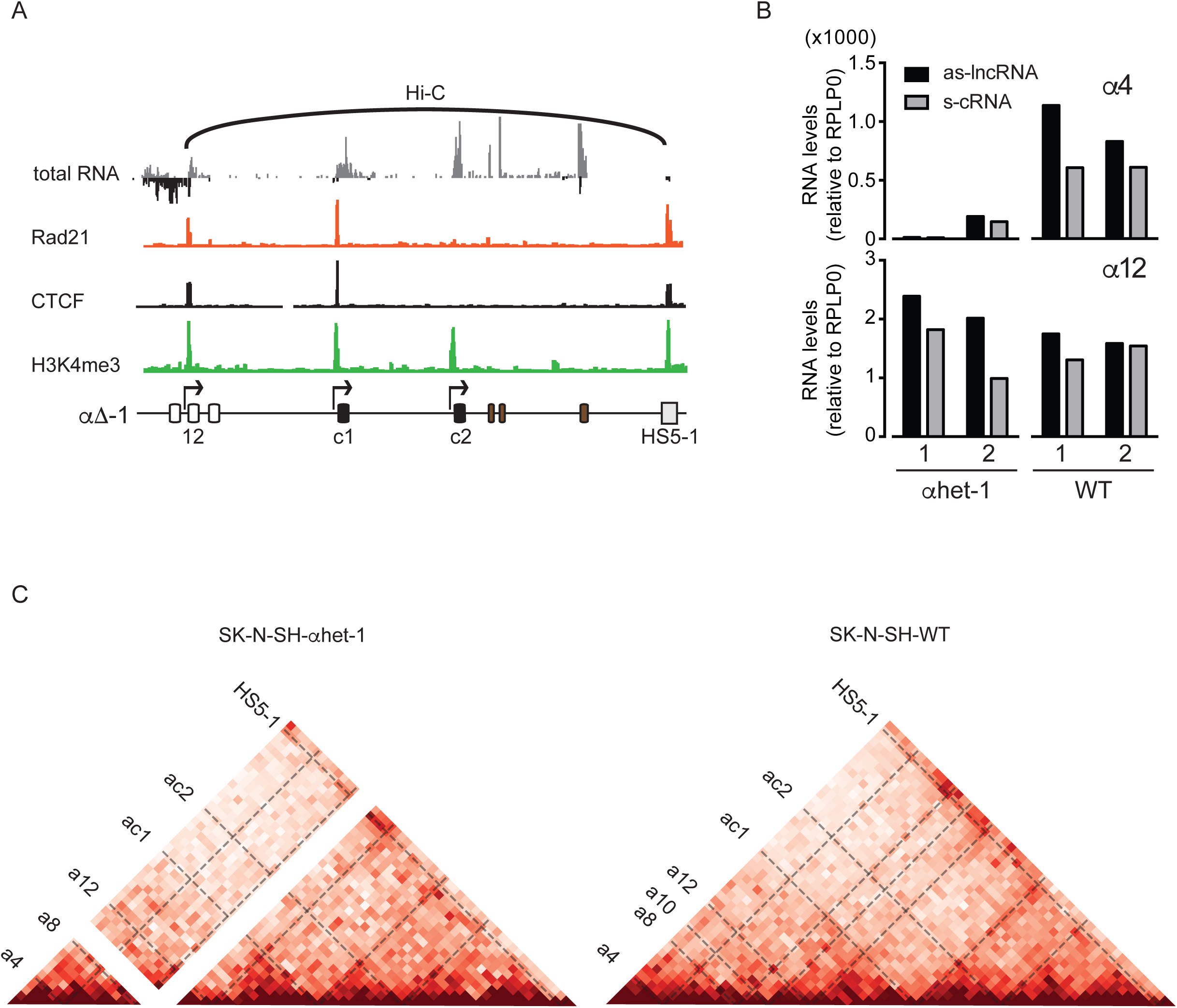
Antisense and sense RNA are transcribed from the same Pcdhα allele. (A) RNA-Seq, ChIP-Seq and *in situ* cHi-C from the SK-N-SHαhet-1. (B) Expression of Pcdhα 4 and 12 in SK-N-SH-αhet 1 and 2 clonal cells compared to SK-N-SH-WT cells. (C) *In situ* cHi-C contact maps at 10kb resolution SK-N-SH-αhet-1 (Left) and SK-N-SH-WT (Right) cells. Coordinates: chr5:140,780,000-141,050,000, hg38.

**Figure S4:**
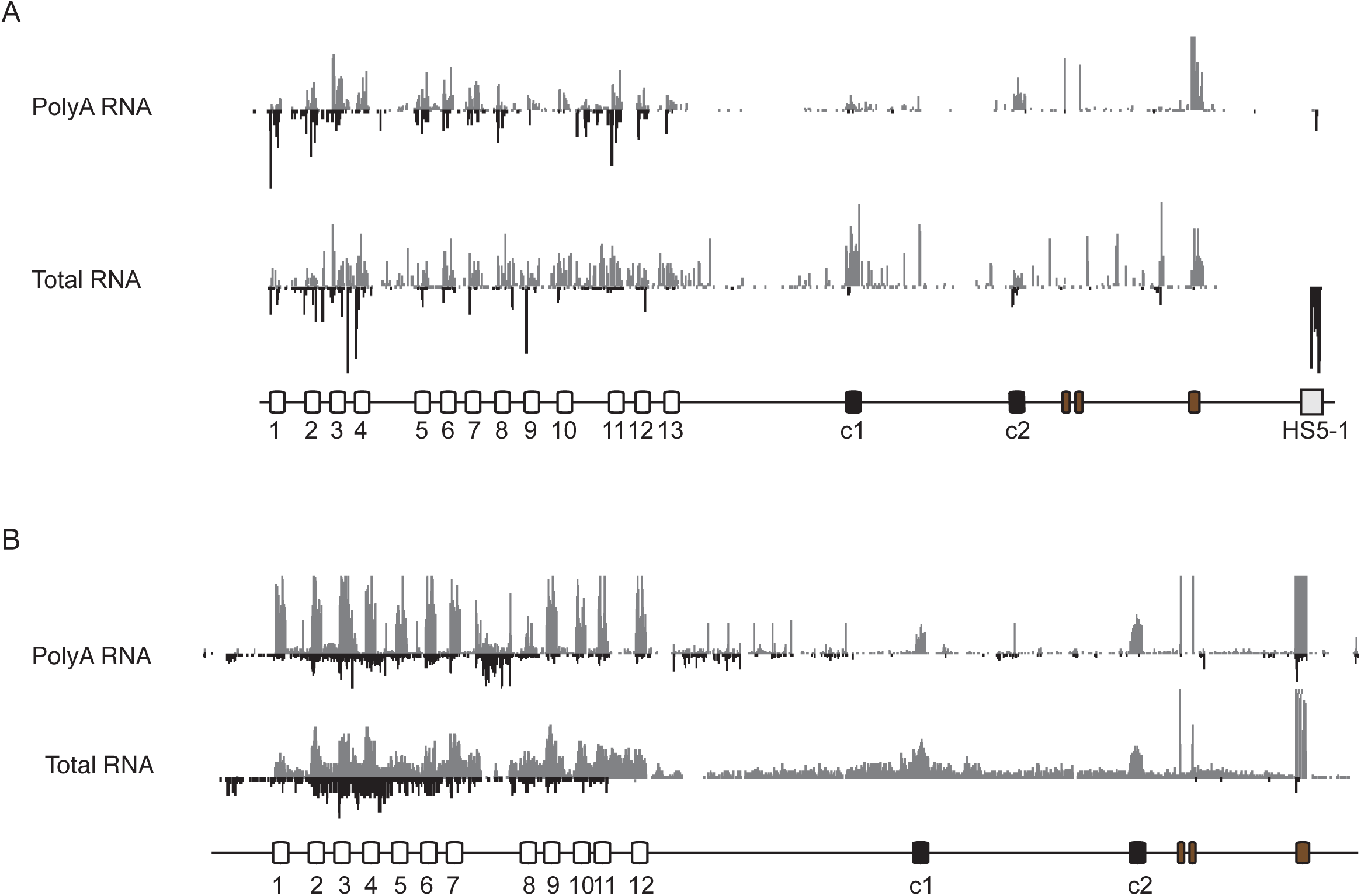
Expression of the Pcdhα cluster *in vivo*. (A) Polyadenylated (PolyA) RNA and Total RNA from human primary neurons. (B) Polyadenylated (PolyA) RNA and Total RNA from mouse olfactory sensory neurons.

**Figure S5:**
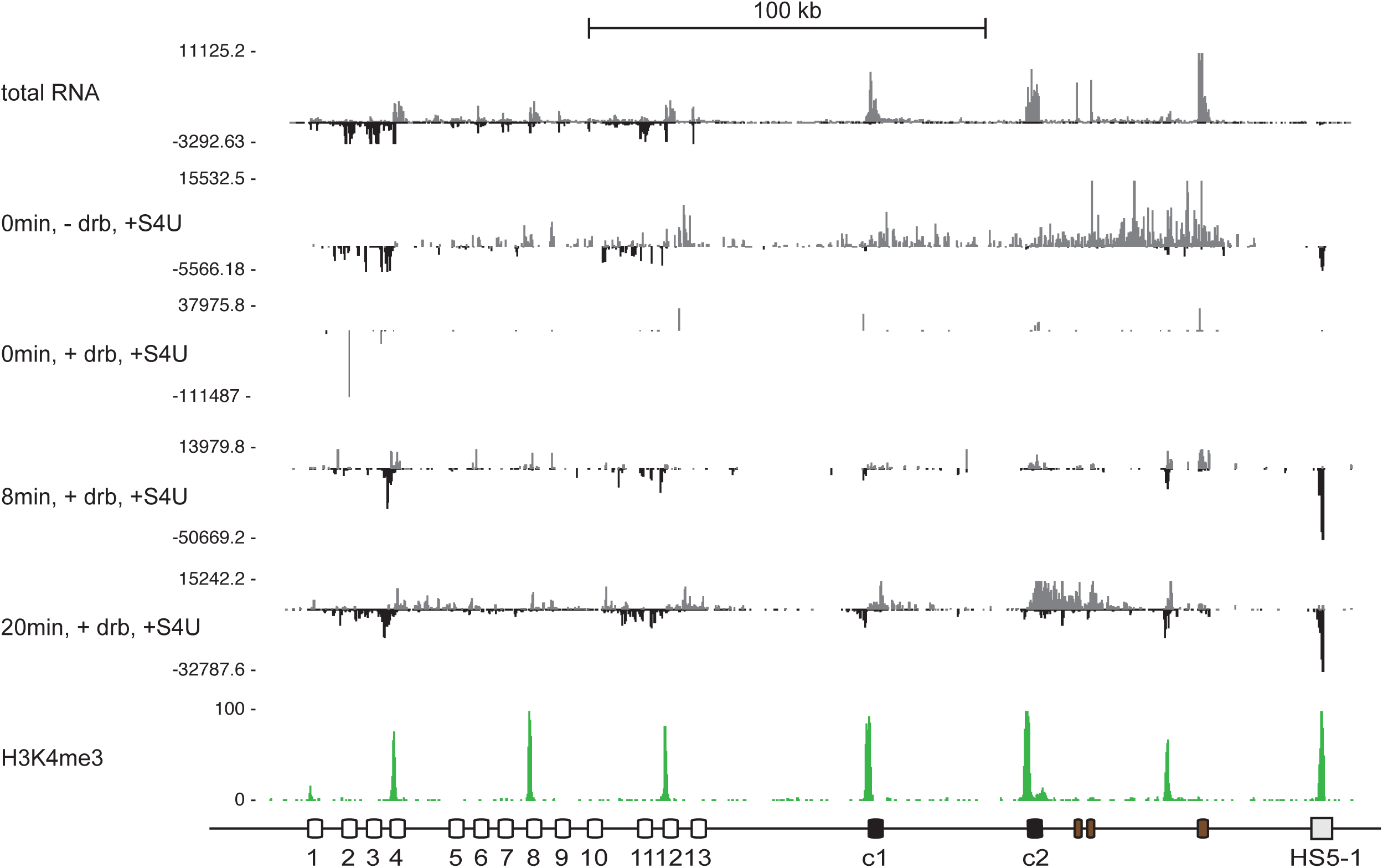
4sUDRB-Seq studies in SK-N-SH cells. Different time points and conditions of s^4^UDRB-Seq relative to total RNA and H3K4me3 in SK-N-SH cells.

**Figure S6:**
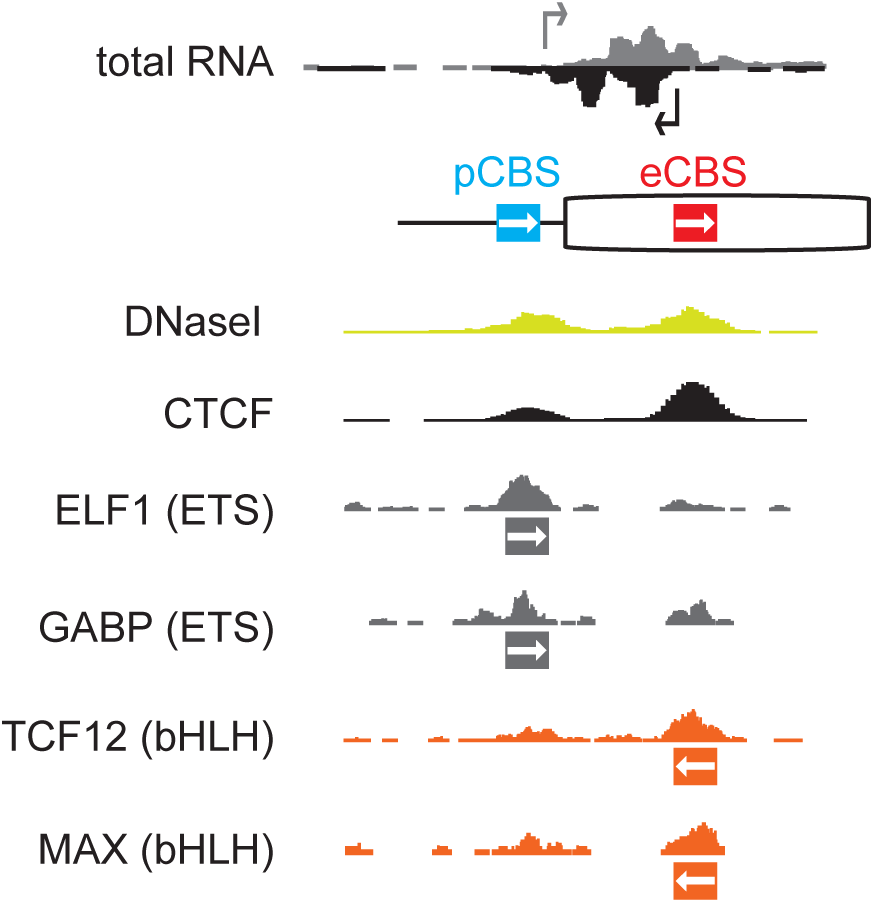
Transcription Factors associated with the Pcdhα convergent promoters. DNaseI hypersensitivity and ChiP-Seq data for distinct transcription factors associated with the active exons in SK-N-SH cells relative to CTCF and total RNA.

**Figure S7:**
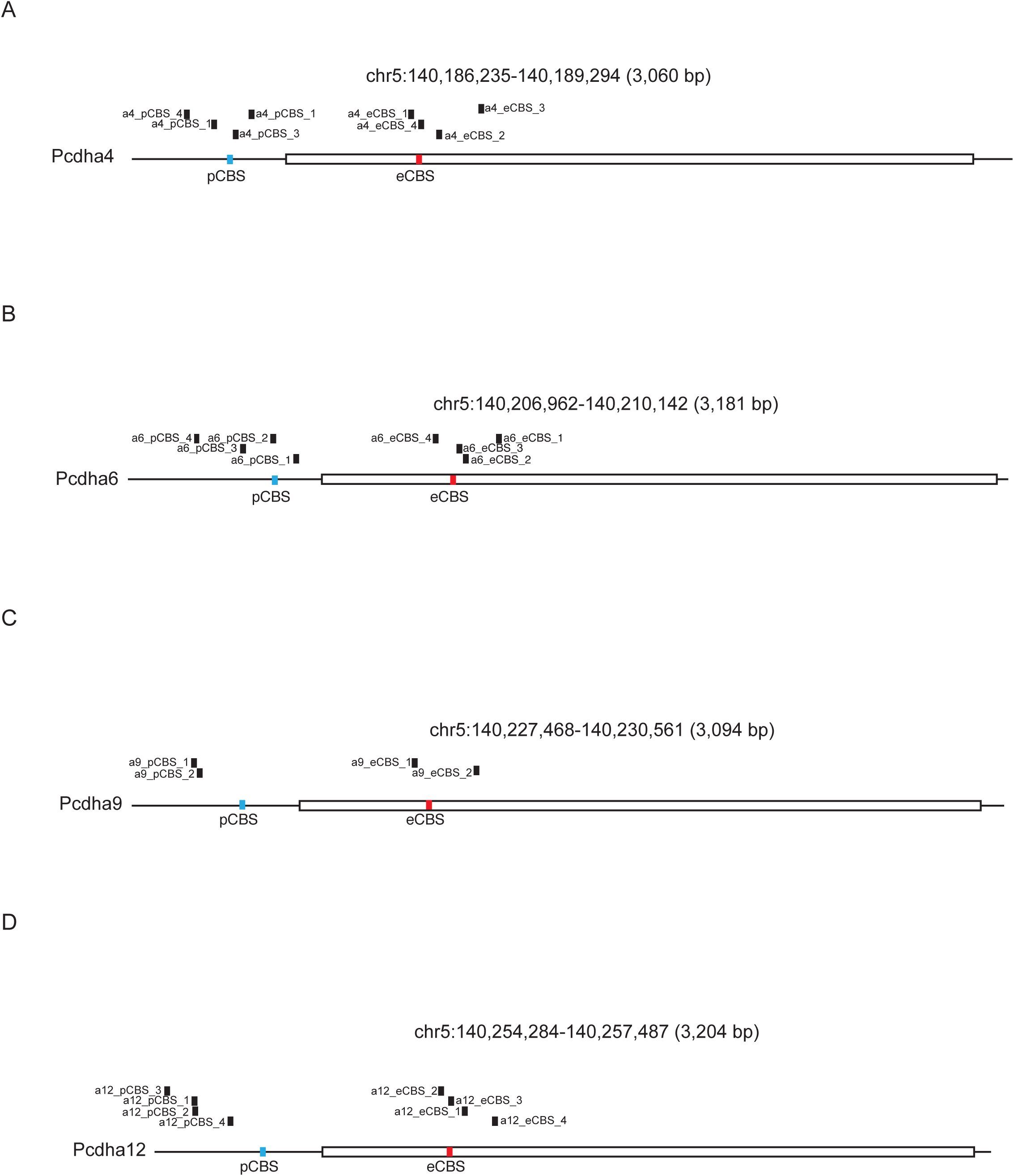
Location of the gRNAs used to activate Pcdhα promoters by dCas9-VPR. Location relative to the pCBS and the eCBS sites of the gRNAs used to activate Pcdh α4 (A), α6 (B), α9 (C) and α12 (D)

**Figure S8:**
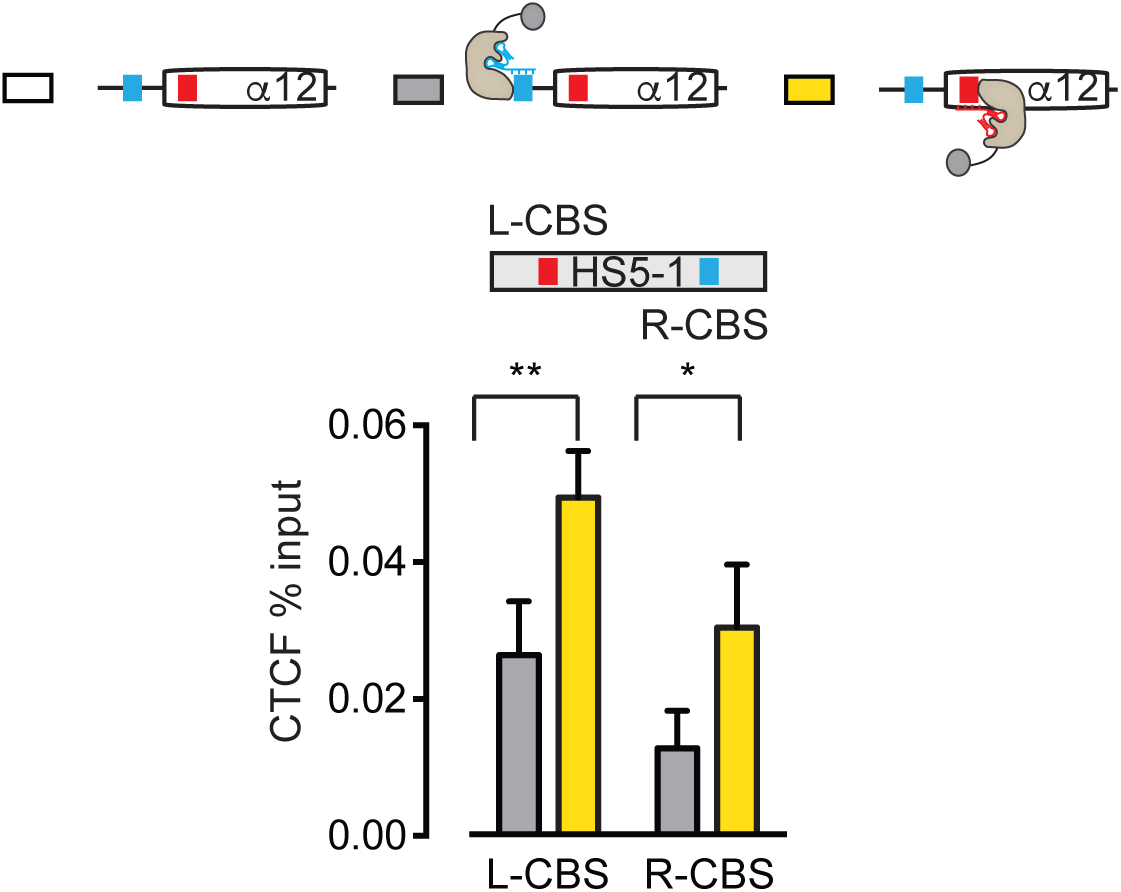
Increase binding of CTCF at the HS5-1 enhancer upon activation of the Pcdhα12 antisense promoter. Enrichment of CTCF occupancy at the L-CBS and the R-CBS sites of the HS5-1 enhancer assayed by ChIP-qPCR. Errors (n>3) represent s.e.m. and statistical significance was calculated with a Student unpaired *t*-test.

**Figure S9:**
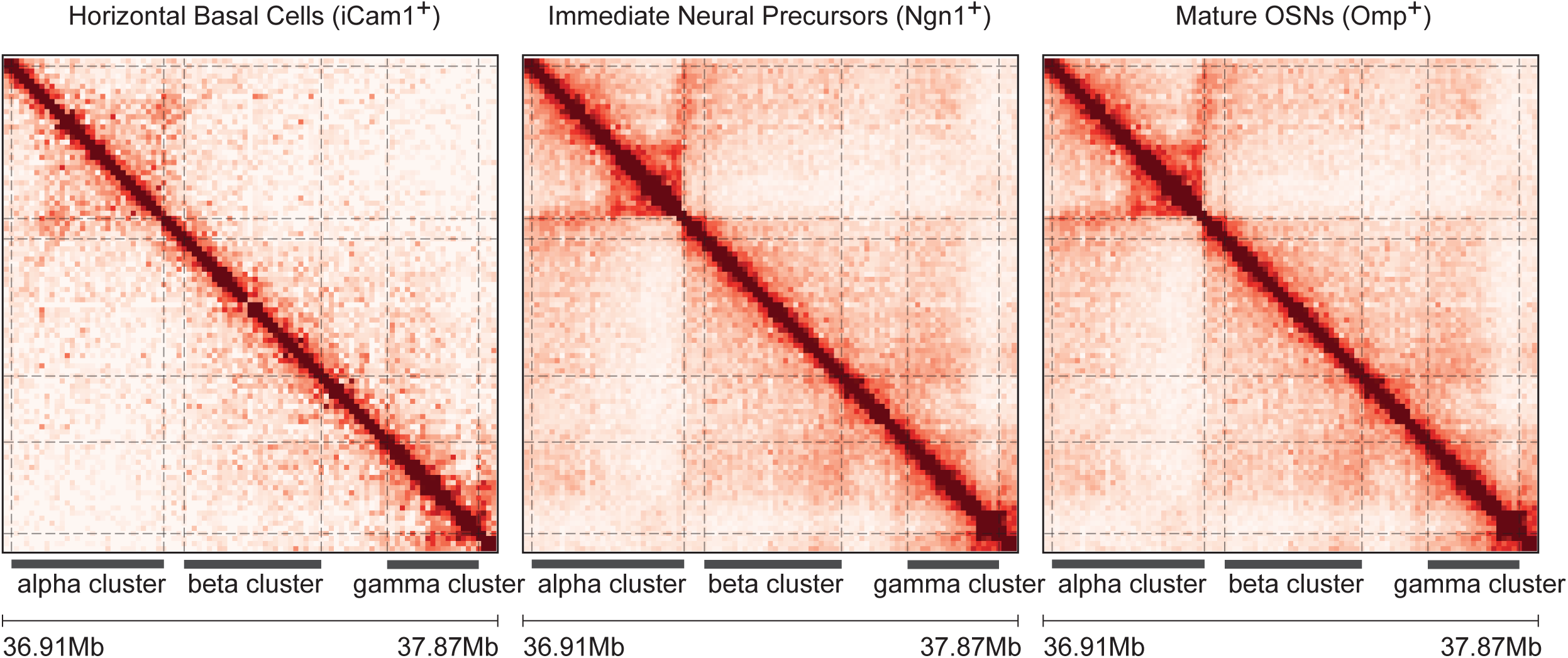
Chromosome architecture of the Pcdh gene cluster during the development of the olfactory epithelium. *In situ* cHi-C contact maps at 10kb resolution for horizontal basal cells (iCam1^+^), immediate neural precursors (Ngn1^+^) and mature olfactory sensory neurons (Omp^+^).

## STAR METHODS

### CONTACT FOR REAGENT AND RESORCE SHARING

Further information and request for resources and reagents should be directed to and will be fulfilled by the Lead Contact, Tom Maniatis.

### EXPERIMENTAL MODEL AND SUBJECT DETAILS

#### Cell lines and Cell culture

SK-N-SH cells were purchased from ATCC and cultured in RPMI-1640 supplemented with 10% (vol/vol) FBS, 1X GlutaMax, 1mM sodium pyruvate, 1X non-essential amino acids, 1% penicillin-streptomycin. HEK293T cells were purchased from ATCC and cultured in DMEM supplemented with 10% (vol/vol) FBS, 1X GlutaMax, 1mM sodium pyruvate, 1X non-essential amino acids, 1% penicillin-streptomycin. Cells were maintained at 37°C in a 5% (vol/vol) CO_2_ incubator.

#### Generation of a CRISPR-inducible SK-N-SH cell line (SK-N-SH-iCas9)

CRISPR-inducible SK-N-SH cells were generated as previously described for Human pluripotent stem cells (hPSCs) (González et al., 2014; Zhu et al., 2014) with the following differences: (1) the Puro-Cas9 donor plasmid was substituted with a GFP-Cas9 donor plasmid and (2) the Neo-M2rtTA donor plasmid was substituted with a mCherry-M2rtTA donor plasmid. Dual color cells were sorted by flow cytometry and genotyped by PCR and further karyotyped.

#### Generation of SK-N-SH heterozygous for the Pcdhα cluster (SK-N-SH-αhet)

SK-N-SH-iCas9 cells were plated at 50% density in a 6-well dish, dox-induced (at a concentration of 2 mg/mL) for 48 hours (refresh Media with 1X RPMI with Dox for every day of induction). On days 3 and 5, the cells were transfected with 1 μg (total) of sgRNAs. On day 6, the GFP/mCherry positive and DAPI negative were single-cell sorted on plates pre-coated with MEF feeder cells. The cells were allowed to grow for a month until visible colonies were observed, replica plated and genotyped by PCR. We isolated two clones (1 and 2) and named this cell line as SK-N-SH-αhet. Deletion of one copy of the Pcdhα cluster in the SK-N-SH-αhet1 clone was further confirmed by a CLG microarray test from Cell Line Genetics (www.clgenetics.com).

#### Animals

Mice were treated in compliance with the rules and regulations of IACUC under protocol number AC-AAAO3902. All experiments were performed on primary FACS-sorted cells from dissected main olfactory epithelium. HBC cells were sorted from keratin5-creER;rt-gfp mice, INP cells were sorted from the brightest GFP populations of ngn1-GFP mice, OSNs were sorted from omp-IRES-GFP mice (Shykind et al., 2004).

### METHODS DETAILS

#### Fluorescence activated cell sorting of HBCs, INPs and OSNs

Cells were dissociated into a single-cell suspension by incubating freshly dissected main olfactory epithelium with papain for 40 minutes at 37°C according to the Worthington Papain Dissociation System. Following dissociation and filtering for three times through a 35 μm cell strainer, cells were resuspended in 1X PBS with 5% FBS. For *in situ* Hi-C experiments, after dissociation, cells were fixed with 1% formaldehyde for 10 minutes at room temperature. Formaldehyde was quenched by adding glycine to a final concentration of 0.125 M for 5 minutes at room temperature. Cells were then washed with 1X cold PBS and resuspended in 1X PBS with 5% FBS. Fluorescent cells were then sorted on a BD Aria II or Influx cell sorter.

#### Transfections of plasmids into HEK293T cells

One day prior to lipid-mediated transfection, HEK293T cells were seeded in a 6-well plate at a density of 1.8^6^ cells per well. For plasmid DNA transfections, 3 μg of total DNA was added to 125 μL of Opti-MEM containing 5 μL of P300 reagent, followed by an addition 125 μL of Opti-MEM containing 7.5 μL of Lipofectamine 3000 per well. The two solutions were mixed and incubated at room temperature for 5 minutes and the solution added dropwise to cells. Plates were then incubated at 37°C for 48 or 72 hours in a 5% CO_2_ incubator. After incubation, cells were harvested in 1 mL of TRIzol.

#### RNA isolation and sequencing

RNA was isolated using TRIzol. Cell lysate was extracted with bromo-chloropropane and RNA was precipitated with 100% isopropanol supplemented with 10 μg of glycoblue for 10 min at room temperature and then pelleted at 16,000 × g for 30 min at 4C. The RNA pellet was washed once with 75% ethanol and then resuspended in RNase-free water to a maximal concentration of 200ng/μl. Genomic DNA contaminants were removed by Turbo DNase. Removal of Turbo DNase was performed by phenol:chloroform extraction and RNA was precipitated as described above and resuspended in RNase-free water and stored at −80C. Sequencing libraries for total RNA and polyadenylated RNA from SK-N-SH cells and human neurons were made using the NEBNext Ultra II Directional RNA Library Prep Kit. Sequencing libraries for total RNA from HEK293T cells and the SK-N-SH-αhet clones were made using the SMARTer Stranded Total RNA-Seq Pico input mammalian RNA kit. The quality of all the libraries was assessed by bioanalyzer and quantified using a combination of bioanalyzer and qubit. Libraries were sequenced on a NEXT-Seq 500/550 at the Columbia Genome Center.

#### Design of the myBaits Capture Library

To overcome the low level of Pcdh expression in both primary neurons and SK-N-SH cells, we made use of an RNA-based enrichment strategy to capture pre-processed and mature RNA species. We refer to this approach as Capture RNA-Sequencing (cRNA-Seq) (see also Figure S1 for a schematic of the myBaits enrichment procedure). myBaits targeted capture kits were designed and purchase from MYcroarray (Arbor Biosciences, http://www.arborbiosci.com). A total of 16,357 biotinylated RNA probes covering about 90.42% of the Pcdh α (chr5: 140159476-140429082 and γ (chr5:140705658-140911381) clusters were synthesized. Baits were design satisfying at least one of the following conditions:

– No blast hit with a T_m_ above 60°C
– No more than 2 hits at 62.5-65°C or 10 hits in the same interval and at least one neighbor candidate being rejected
– No more than 2 hits at 65-67.5°C and 10 hits at 62.5-65°C and two neighbor candidates on at least one side being rejected
– No more than a single hit at or above 70°C and no more than 1 hit at 65-67.5°C and 2 hits at 62.5-65°C and two neighbor candidates on at least one side being rejected

Sequencing libraries from RNA-Seq or HiC-Seq were multiplexed at the desired ratio and captured using the myBaits Capture Library protocol for 18 hours at 65°C. Captured libraries were eluted in RNase-free water and further amplified. The quality of captured libraries was assessed by bioanalyzer and quantified using a combination of bioanalyzer and qubit. Libraries were sequenced on a NEXT-Seq 500/550 at the Columbia Genome Center.

#### RNAPII pausing

Start-Seq experiments were previously described Nechaev et al., 2010) with the following changes: (1) about 10^6^ SK-N-SH cells were used for each replicate experiment and (2) RNA-Seq libraries were prepared with the NEXTflex small RNA kit v3. The 3’end of the Start-RNA libraries were sequenced to determine the location of RNAPII pausing.

#### RNAPII elongation

SK-N-SH cells were treated with 100 μM DRB or DMSO for 6 hours. s^4^UDRB experiments were performed as previously described (Fuchs et al., 2014; 2015) with the following changes: 1 mM s^4^U was added to media 20 min before cells were harvested. After 6h, DRB and s^4^U-containing media was removed and replaced with s^4^U-containing media, and cells were harvested with TRIzol after 0, 8, or 20 min after DRB removal. Cells were flash frozen and stored at −80°C. A no DRB and a no s^4^U controls were also performed.

Total RNA was purified and s^4^U-RNA was enriched using MTS-biotin chemistry (Duffy and Simon, 2009; Duffy et al., 2015). Briefly, cells were lysed in TRIzol, extracted with chloroform once and the nucleic acids precipitated with isopropanol. DNA was removed with Turbo DNase. DNase protein was removed by phenol:chloroform:isoamylalcohol extraction, and RNA isolated using isopropanol precipitation. RNA was sheared to ~200 bp by adding shearing buffer (150mM Tris-HCl pH 8.3, 225mM KCl, 9mM MgCl_2_) and heating to 94°C for 4min, followed by quenching on ice with EDTA. Sheared RNA was purified using a modified protocol with the RNeasy Mini Kit (Qiagen). To biotinylate the s^4^U-RNA, 150μg sheared RNA was incubated with 60μg MTS-biotin in biotinylation buffer (150 μL total volume) for 30min. Excess biotin was removed via chloroform extraction using Phase-Lock Gel Tubes. RNA was precipitated with a 1:10 volume of 3M NaOAc and an equal volume of isopropanol and centrifuged at 20,000 × g for 20min. The pellet was washed with an equal volume of 75% ethanol. Purified RNA was dissolved in 200μl RNase-free water. Biotinylated RNA was separated from non-labelled RNA using glycogen-blocked Dynabeads Streptavidin C1 Beads (Invitrogen). Beads (200μl) were added to each sample and incubated for 15min at room temperature, then washed three times with high salt wash buffer (1 ml each, 100mM Tris-HCl (pH 7.4), 10mM EDTA, 1M NaCl, and 0.1% Tween-20). In order to improve the stringency of the washes, an additional three washes with buffer TE (10mM Tris pH 7.4, 1mM EDTA) at 55°C were performed. s^4^U-RNA was eluted from Dynabeads with 200μl freshly prepared elution buffer (10mM DTT, 100mM NaCl, 10mM Tris pH 7.4, 1mM EDTA) and incubated for 15min. Enriched RNA was purified by ethanol precipitation and re-biotinylated as above. Excess biotin was removed via chloroform extraction using Phase-Lock Gel Tubes and RNA was purified by RNeasy Mini Kit. s^4^U-RNA was enriched on streptavidin beads as above and beads were washed three times with high salt wash buffer. s^4^U-RNA was eluted as above and spiked with 200 pg *Schizosaccharomyces pombe* total RNA. 10 ng total RNA from input and enriched RNA samples was used for library preparation with the SMARTer Stranded Total RNA-seq Kit Pico Input Mammalian (Clontech) according to the manufacturer’s instructions. Input and enriched samples were multiplexed with Illumina barcodes and sequenced using paired-end 2×75-nt cycles on an Illumina NextSeq 500/550 instrument.

#### Chromatin Immunoprecipitation (ChIP-Seq and ChIP-qPCR)

The following antibodies were used for chromatin immunoprecipitation studies: CTCF (donated by Victor Lobanenkov), Rad21 (Abcam ab992), Histone H3 Lysine 4 tri-methyl (ThermoFisher PA5-27029), Histone H3 Lysine 27 acetylation (Abcam ab4729), FLAG (Sigma F1804). Cells were crosslinked with 1% formaldehyde for 10 minutes at room temperature. Formaldehyde was quenched by adding glycine to a final concentration of 0.125 M for 5 minutes at room temperature. Cells were then washed with 1X cold PBS with protein inhibitors twice and pelleted. Cell pellets were stored at −80C till use. Cells were lysed in lysis buffer (50 mM Tris pH 7.5, 140 mM NaCl, 0.1% SDS, 0.1% sodium deoxycholate, 1% Triton X-100) for 10 minutes. Nuclei were span for 10 minutes at 1000g and resuspended in the sonication buffer (10 mM Tris pH 7.5, 0.5% SDS) as 5^6^ nuclei per 300 μl sonication buffer. Chromatin was sheared by bioruptor for 30 cycles at cycle condition 30/30 (ON/OFF time in seconds). Following a spin at 13,000g for 10 minutes to remove debris, the sheared chromatin was diluted such as the final binding buffer concentration was 15 mM Tris-HCl pH 7.5, 150 mM NaCl, 1 mM EDTA, 1% Triton X-100, 0.1% SDS) and incubated for 2 hours with dynabeads G pre-equilibrated in the binding buffer for pre-clearing of the chromatin. Post-cleared chromatin was then incubated with the specific antibody overnight (1 μg of antibody was used per 5^6^ nuclei). The next day, dynabeads G were added to the chromatin-antibody mix for 2 hours. A total of 4 washes with 1X wash buffer (100 mM Tris pH 7.5, 500 mM LiCl, 1% NP-40, 1% sodium deoxycholate) and 1 wash with TE buffer (10 mM Tris pH 7.5, 1 mM EDTA) were performed. The elution was performed at 65°C for 1 hour in the elution buffer (1% SDS, 250 mM NaCl, 2 mM DTT). All steps, with the exception of the elution, were performed at 4°C. All buffers, with the exception of the TE and elution buffer contained 1X protease inhibitors. The eluted chromatin was reverse-crosslinked overnight at 65°C and the DNA was purified with the Zymo DNA kit.

Libraries for ChIP-Seq were prepared using the NEBNext Ultra II DNA Library Prep Kit. The quality of the libraries was assessed by bioanalyzer and quantified using a combination of bioanalyzer and qubit. Libraries were sequenced on a NEXT-Seq 500/550.

#### *In situ* Chromatin Capture Conformation (Hi-C)

1.5^6^ SK-N-SH or HEK293T cells were lysed and intact nuclei were processed through an *in situ* Hi-C protocol as previously described with a few modifications (Rao et al., 2014). Briefly, cells were lysed with 50 mM Tris pH 7.5 0.5% Igepal, 0.25% Sodium-deoxychloate, 0.1% SDS, 150 mM NaCl, and protease inhibitors. Pelleted intact nuclei were then resuspended in 0.5% SDS and incubated for 20 minutes at 65°C for nuclear permeabilization. After quenching with 1.1% Triton-X for 10 minutes at 37°C, nuclei were digested with 6 U/μl of DpnII in 1x DpnII buffer overnight at 37°C. Following initial digestion, a second DpnII digestion was performed at 37°C for 2 hours. DpnII was heat-inactivated at 65°C for 20 minutes. For the 1.5hr fill-in at 37°C, biotinylated dGTP was used instead of dATP to increase ligation efficiency. Ligation was performed at 25°C for 4 hours. Nuclei were then pelleted and sonicated in 10 mM Tris pH 7.5, 1 mM EDTA, 0.25% SDS on a Covaris S220 for 16 minutes with 2% duty cycle, 105 intensity, 200 cycles per burst, 1.8-1.85 W, and max temperature of 6°C. DNA was reverse cross-linked overnight at 65°C with proteinase K and RNAse A. Reverse cross-linked DNA was purified with 2× Ampure beads following the standard protocol. Biotinylated fragments were enriched using Dynabeads MyOne Strepavidin T1 beads. The biotinylated DNA fragments were prepared for next-generation sequencing on the beads by using the Nugen Ovation Ultralow kit protocol with some modifications. Following end repair, magnetic beads were washed twice at 55°C with 0.05% Tween, 1 M NaCl in Tris/EDTA pH 7.5. Residual detergent was removed by washing beads twice in 10 mM Tris pH 7.5. End repair buffers were replenished to original concentrations, but the enzyme and enhancer was omitted before adapter ligation. Following adaptor ligation, beads underwent 5 washes with 0.05% Tween, 1 M NaCl in Tris/EDTA pH 7.5 at 55°C and two washes with 10mM Tris pH 7.5. DNA was amplified by 10 cycles of PCR, irrespective of starting material. Beads were reclaimed and amplified unbiotinylated DNA fragments were purified with 0.8x Ampure beads. Quality and concentration of libraries were assessed by Agilent Bioanalyzer and Qubit. *In situ* Hi-C libraries from SK-N-SH and HEK293T cells were size-selected and enriched as described above using the myBaits Capture Library protocol described above and sequenced paired-end on NextSeq 500 (2×75bp).

#### Methylated DNA Immunoprecipitation (MedIP)

The following antibodies were used: 5-Methylcytosine (5-mC) antibody (Active Motif 39649) and 5-Hydroxymethylcytosine (5-hmC) antibody (Active Motif 39791).

HEK293T cells were transfected with the appropriate set of dCas9 plasmids and incubated at 37°C for 72 hours in a 5% CO2 incubator. Genomic DNA was extracted using the PureLink Genomic DNA Mini Kit (Invitrogen). A total of 2 μg of DNA was diluted into 300 μl TE sonication buffer (10 mM Tris pH 7.5, 1 mM EDTA). Genomic DNA was sheared by bioruptor for 18 cycles at cycle condition 30/90 (ON/OFF time in seconds). The sheared DNA was diluted to a final IP buffer of 15 mM Tris-HCl pH 7.5, 150 mM NaCl, 1 mM EDTA, 1% Triton X-100 and incubated overnight with 1 μg of antibody. The next day, a mixture of dynabeads A and G were added to the DNA-antibody mix for 2 hours. A total of 3 washes with 1X IP buffer were performed. The elution was performed at 55°C for 3 hour with rigorous shaking in the elution buffer (1% SDS, 250 mM NaCl). All steps, with the exception of the elution, were performed at 4°C. The eluted DNA was purified with the Zymo DNA kit.

#### Bioinformatic Analysis of Sequencing Data

For RNA-Seq experiments, raw FASTQ files were aligned with either Tophat or STAR using hg19 or mm10 reference genomes. When libraries were made following the SMARTer Stranded Total RNA-Seq, the initial 4 base pairs of both paired reads were trimmed prior to alignment.

For ChiP-Seq experiments, raw FASTQ files were aligned using Bowtie2 using hg19 reference genome upon adapter sequences removal using CutAdapt. Uniquely aligning reads were selected using Samtools and reads with alignment quality below 30 (-q 30) were removed. The HOMER software package was used to generate signal tracks.

For *in situ* Hi-C experiments, raw FASTQ files were processed through use of the Juicer Tools Version 1.76 pipeline (Durand et al., 2016) with one modification. Reads were aligned to hg38 using BWA 0.7.17 mem algorithm and specifying the −5 option implemented specifically for *in situ* Hi-C data. For captured Hi-C libraries, contact matrices were normalized to 2kb resolution by first reporting counts as reads per billion Hi-C contacts, then by normalizing with the Knight Ruiz (KR) matrix balancing algorithm (Knight and Ruiz, 2013) focused on the alpha Pcdh cluster (chr5:140780000-141046000; hg38). For uncaptured libraries (mm10 Hi-C), matrices were KR normalized genome wide.

For generating a contact matrix, scales were set to a minimum of 0 reads and a maximum of 2^∗^(mean normalized reads) in order to report a relative enrichment of contacts.

Local peak analysis was performed by normalizing contacts between HS5-1 (chr5: 141020000-141066000) and alternative Pcdh exons (chr5:140850000-140898000) to reads per billion Hi-C contacts at 5kb resolution. Scales were set to 0 and the maximum value for each plot.

DNaseI and ChIP data for H3K4me3, CTCF, Rad21, ELF1, GABP, TCF12, MAX, YY1 in SK-N-SH cells were obtained from the ENCODE data matrix.

*In situ* Hi-C data for INP and OSN cells were obtained from (Horta et al., 2018).

#### CRISPR gRNA design

All guide RNA (gRNAs) were designed as truncated 18mer long sequences to increase their binding specificity as previously described (Fu et al., 2014) using the CRISPR design web tool (http://crispr.mit.edu). With the exception of the Pcdhα9, where a total of 2 gRNAs were used for activate either the pCBS- or the eCBS-promoters, we used 4 gRNAs for the activation of the pCBS- and eCBS-promoters of Pcdh α4, α6, α12.

#### *In vitro* transcription of gRNAs

The gRNAs were transcribed using the MEGAshortscript T7 Transcription Kit by Life Technologies (AM1354M), purified by phenol-chloroform and transfected in the SK-N-SH-iCas9 cells by RNAimax lipofectamine reagent.

### QUANTIFICATION AND STATISTICS

The statistical tests used in this study are indicated in the respective figure legends. In general, data with single independent experiments were analyzed by Student unpaired *t*-test to determine statistical significant effects (p < 0.05). Data with multiple independent experiments were analyzed by one-way ANOVA to determine statistical significant effects (p < 0.05).

### DATA AND SOFTWARE AVAILABILITY

The data discussed in this work will be available upon request.

## SUPPLEMENTAL DATA TABLES

**Supplemental Data Table 1:** Table of the primers used in this study

| REAGENT or RESOURCE | SOURCE | IDENTIFIER |
| --- | --- | --- |
| Antibodies |  |  |
| CTCF monoclonal | Pugacheva et al, 2015 | N/A |
| Rabbit polyclonal anti Rad21 | Sigma-Aldrich | ab922 |
| Rabbit polyclonal anti H3K4me3 | ThermoFisher | PA5-27029 |
| Rabbit polyclonal anti H3K27ac | Abcam | ab4729 |
| Mouse monoclonal anti Flag | Sigma | F1804 |
| Biological Samples |  |  |
| Fetal brain neurons | Dr. Ben Barres | N/A |
| Chemicals, Peptides, and Recombinant Proteins |  |  |
| 5,6-Dichloro-1-β-D-ribofuranosylbenzimidazole (DRB) | Sigma | D1916-50MG |
| 4-Thiouridine, 98+% | Alfa Aesar | J60679 |
| Critical Commercial Assays |  |  |
| SMARTer® Stranded Total RNA-Seq Kit - Pico Input Mammalian | Clontech | 635005 |
| NEBNext® Ultra™ II DNA Library Prep Kit for Illumina | NEB | E7645S |
| Deposited Data |  |  |
| Raw and analyzed data | This paper | GEO: X |
| Experimental Models: Cell Lines |  |  |
| SK-N-SH cells | ATCC | HTB-11 |
| HEK293T cells | ATCC | CRL-1573 |
| Experimental Models: Organisms/Strains |  |  |
| Mouse: R6/2: B6CBA-Tg(HDexon1)62Gpb/3J | The Jackson Laboratory | JAX: 006494 |
| keratin5-creER;rt-gfp mice | Lomvardas Lab | N/A |
| ngn1-GFP | Lomvardas Lab | N/A |
| omp-IRES-GFP | Lomvardas Lab | N/A |
| Recombinant DNA |  |  |
| CMV-dCas9-VPR | This paper | N/A |

**Supplemental Data Table 2:**
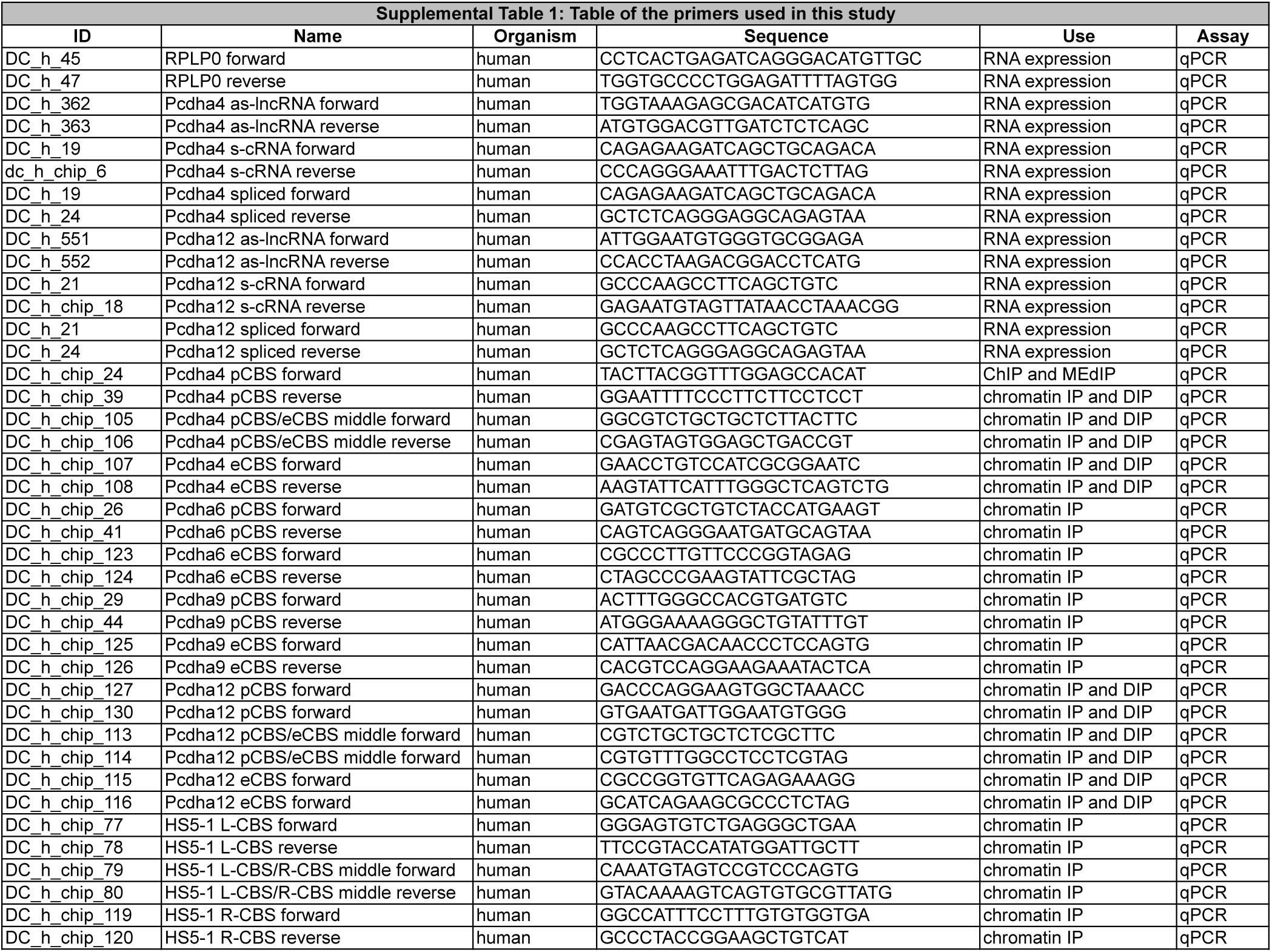
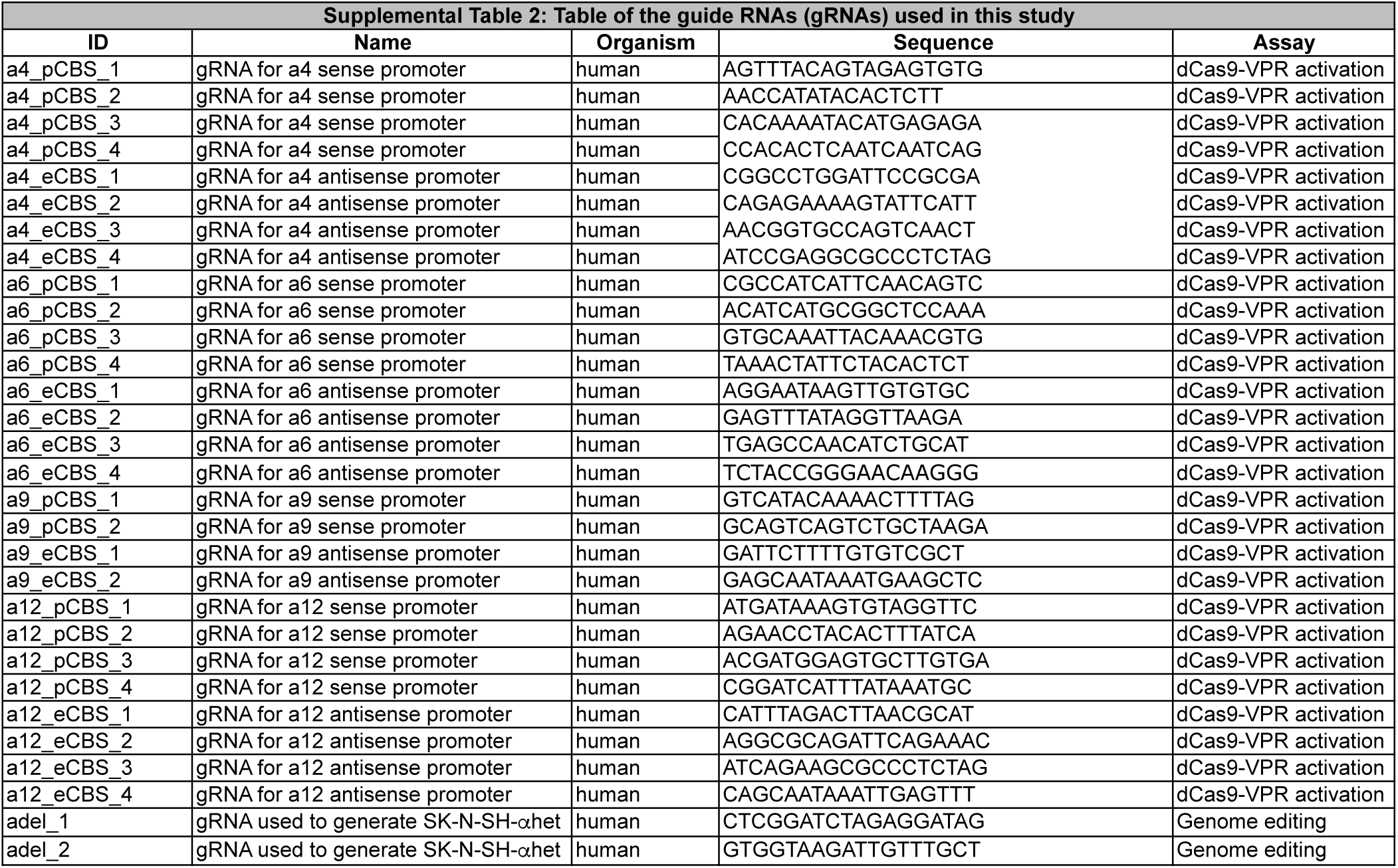
Table of the guide RNAs (gRNAs) used in this study

